# The role of structural *vs* cellular remodeling in arrhythmogenesis: personalized computer models of atrial fibrillation

**DOI:** 10.1101/2023.05.13.540632

**Authors:** Andrey V. Pikunov, Roman A. Syunyaev, Rheeda Ali, Adityo Prakosa, Patrick M. Boyle, Vanessa Steckmeister, Ingo Kutschka, Eric Rytkin, Niels Voigt, Natalia Trayanova, Igor R. Efimov

**Author notes:** – Corresponding authors. Email:,. – These authors contributed equally to this work (Andrey V. Pikunov, Roman A. Syunyaev).

## Abstract

Atrial fibrillation (AF) is a progressive disease involving both structural and functional remodeling. To investigate the contribution of cell-scale functional remodeling to AF pathogenesis, we combined personalized 3D anatomical models with pathology-specific ionic models. The latter were developed using recordings in myocytes isolated from patients in sinus rhythm, paroxysmal, postoperative, and persistent AF. To quantify AF dynamics, we developed a novel algorithm for locating RDs by backtracking the conduction velocity field from the wavebreak regions. We demonstrate that our novel algorithm is at least 700 times faster than the traditional phase singularity analysis. The inducibility of simulated AF was not pathology-dependent, but pathological models demonstrate a more extensive arrhythmogenic substrate compared to the sinus rhythm. AF driver locations depend on electrophysiological remodeling; differences between pathology-specific models are explained by differences in wavebreak patterns. Specifically, RDs tend to dwell in the regions with the highest wavebreak probability.

## Introduction

Atrial fibrillation (AF) is the most common cardiac arrhythmia worldwide (*1*). AF, by its nature, is a progressive disease (*2, 3*), initial episodes are most usually self-terminating with episodes lasting less than 7 days and classified as paroxysmal AF (pAF). Over time it progresses towards persistent AF (persAF), with episodes longer than 7 days, long-standing AF (>6 months), and permanent AF. Another presentation of AF is postoperative AF (poAF), which occurs as a complication in 30-40% of patients following cardiac surgery (*4–6*). Two mechanisms responsible for sustaining arrhythmogenic activity in AF are focal ectopic activity and re-entrant drivers (*7*). The AF episode initiation requires two conditions: an initiating trigger, such as an ectopic excitation triggered by a delayed afterdepolarization, and arrhythmogenic substrate (*8*). The current understanding of AF pathophysiology is formulated as “’AF begets AF”, that is, AF episodes promote the arrhythmogenic substrate via structural and electrophysiological remodeling: tissue fibrosis, gap-junctions downregulation, and action potential (AP) shortening (*2, 3*). Despite advances, the management of AF remains suboptimal due to the complexity and diversity in the interaction between mechanisms underlying AF (*4, 5*).

Antiarrhythmic drug treatment of AF currently suffers from limited efficacy and a significant risk of adverse effects (*9*). A more successful strategy of AF treatment, catheter ablation, has emerged over the last few decades (*10–13*), with pulmonary vein (PV) isolation proving particularly effective (*10–12*). Early on, PVs were thought to be a source of focal ectopic activity, however, later studies suggested PVs to be also a preferential region of reentry formation (*8*). While PV isolation was successful in pAF patients, targeting PV was less effective in persAF due to the wider arrhythmogenic substrate (*13, 14*). More recently, computationally guided ablation was proposed (*15, 16*): patient-specific computational models accounting for fibrotic remodeling are reconstructed from late gadolinium-enhanced MRI scans (LGE-MRI), then optimal targets for ablation are determined via biophysically detailed simulations. Although the potential efficacy of this technique was shown in recent studies (*15, 17, 18*), patient-specific remodeling on a myocyte level was not accounted for so far, the same single-cell AF AP model was used for computations. In the current study, we demonstrate the dependence of arrhythmogenic substrate on electrophysiological remodeling at the cellular level. The pipeline we propose here combines patient-specific electrophysiological models reconstructed from patch-clamp AP recordings with patient-specific geometry and fibrotic remodeling reconstructed from LGE-MRI scans (Figure 1A). While myocardial biopsy required for the patch-clamp recordings is, in general, not justified clinically, different techniques such as monophasic AP recordings might be used for model personalization (*19*).

**Figure 1.**
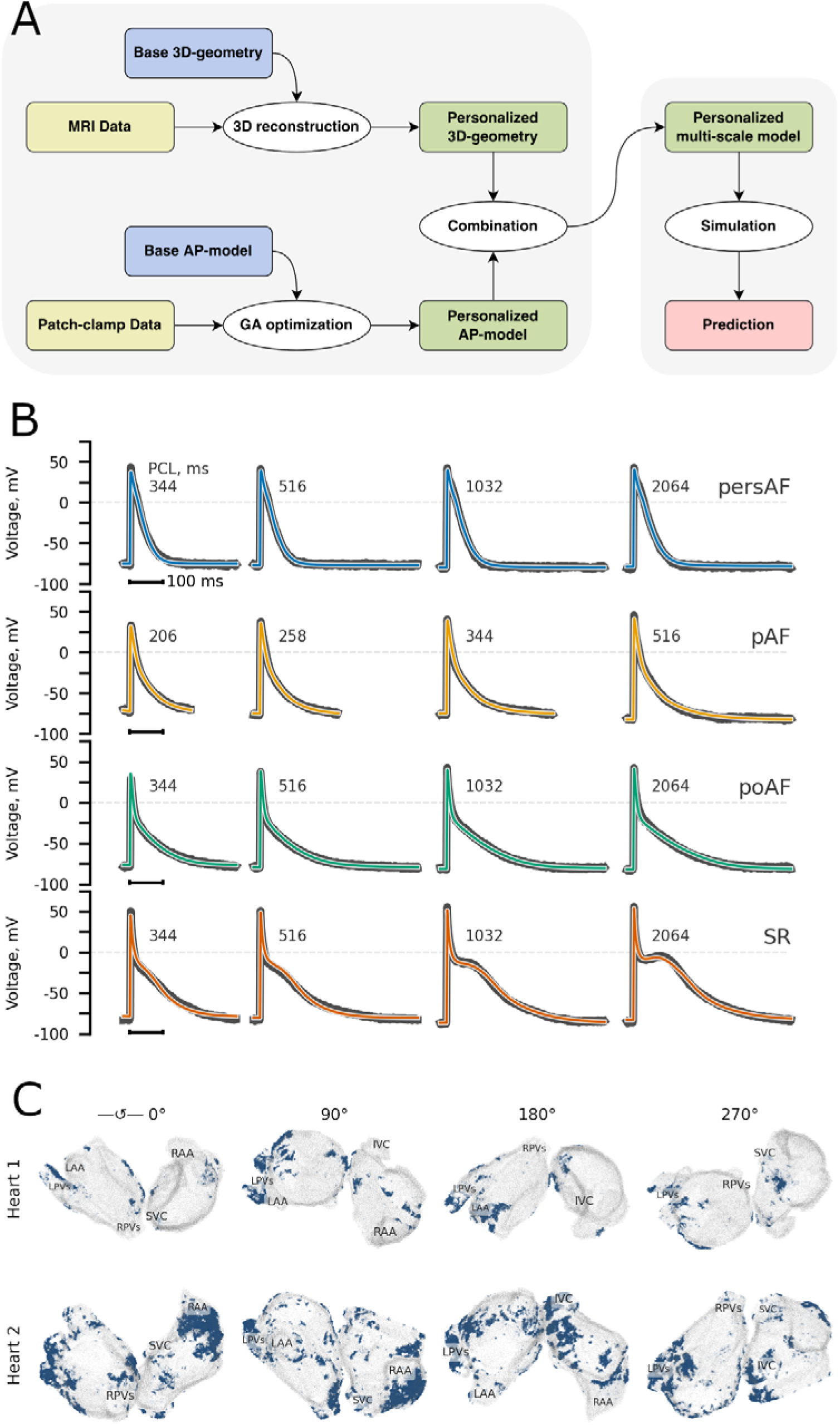
Personalization pipeline. A. Multiscale personalization pipeline combines patch-clamp and late Gd enhanced MRI data. B. We personalized the Maleckar AP model to fit AP waveform restitution measured via the patch-clamp technique. Four traces correspond to four types of patients: persistent AF (persAF), paroxysmal AF (pAF), postoperative AF (poAF), and sinus rhythm (SR). C. Two patient-specific 3D geometries of atria alongside fibrosis distribution were obtained using LG-MRI data. Model 1 and 2 (from now on, mentioned as Heart 1 and 2, respectively) have 10% and 25% fibrosis, respectively. In case of Heart 1, LPVs, LAA, and septum have the highest abundance of fibrotic tissue; in case of Heart 2, LPVs and RAA. The columns of the figure are different views (rotated around the horizontal axis). We use the first view (0°) for the subsequent 3D figures.

Ablation procedures generally require that electrophysiologists determine the locations of reentrant or focal drivers, which is often confusing and time-consuming due to the spatiotemporal instability of AF (*20*). Incorrect identification of the source of the arrhythmogenic activity or suboptimal choice of ablation targets can actually create new substrates for reentrant drivers (RDs) instead of terminating AF (*21*). In non-clinical studies, when not only the activation sequence but actual AP recordings might be available to the researcher, several algorithms exist to automatically detect the RDs. The most widely used is phase singularity (PS) analysis (*22*). However, automatic PS detection is computationally expensive when a researcher intends to process many simulations. Moreover, this technique often results in both false positive and false negative PSs (*23, 24*). Since correct identification of RDs remains a challenging problem both in computational and clinical studies, new algorithms are being developed, such as the algorithms using directed graph networks (*25, 26*). In our study, we propose a novel RD detection algorithm that backtracks the conduction velocity (CV) field to the activation source and then uses a neural network to automatically locate quasi-periodic activation sources. The backtracking algorithm (BTA) uses only the activation sequence as input data. We demonstrate it to be robust and computationally efficient enough to potentially apply to clinical studies based on activation sequences.

## Results

### Model Optimization

In AF, cardiac tissue undergoes both structural and electrophysiological (*44–46*) remodeling: both fibroblast proliferation and changes in myocyte expression profile affect the arrhythmogenesis. In this article, we aimed to investigate the effects of the latter: how changes to the AP waveform and restitution affect arrhythmogenesis given the same structural remodeling. Therefore, we have combined four pathology-specific myocyte models corresponding to SR, poAF, pAF and persAF (**Figure 1B**, see ‘Materials and methods’ for details) with patient-specific geometry based on LGE-MRI of persistent AF patients (**Figure 1C**)(*15*). The pathological myocyte phenotypes were characterized by a somewhat shorter and more triangular AP waveform (**Table S4**) as described previously (*2, 8, 46–48*). SR and three pathological GA-optimized Maleckar models (pAF, persAF, and poAF) reproduced experimental AP waveforms with high precision: Root Mean Square Error (RMSE) between model and experimental AP waveforms was 1.64 ± 0.43 mV (see also **Table S4**). Since both patient-specific geometries we used in the study corresponded to persistent AF patients, they were characterized by extensive fibrotic remodeling, most notably around pulmonary veins (both hearts) and right atrial appendage (Heart 2).

### RD location

In order to automatically locate RDs induced *in silico,* we have developed a novel algorithm (**Figure 2**). Briefly (see ‘Materials and methods’ for details), we backtrack the velocity field to find an excitation source that can be either focal or quasi-periodic RD. A more traditional algorithm widely used in both computational and experimental studies is based on PS location; phase is calculated via Hilbert transform (HT) (*43*). **Figure 3** shows the difference between HT and RD as defined by BTA. The reentry waves rotate around either an anatomical obstacle or an area of functional block (*49*), constituting the core of the reentry (white area in **Figure 3A**). In the latter case, the PS is defined as a site where all of the phases of oscillations converge; in contrast, the surrounding elements exhibit a continuous progression of phase (*50*). As demonstrated in **Figure 3**, the tissue within RD is electrotonically activated, and the PS is actually located within the reentrant core, where myocytes undergo subthreshold oscillations (**Figure 3C**). On the other hand, RD is defined as the actual arrhythmogenic source of excitation. It is also important to note that while voltage recording is required for HT, only the activation sequence is used instead in our algorithm to backtrack the CV.

**Figure 2.**
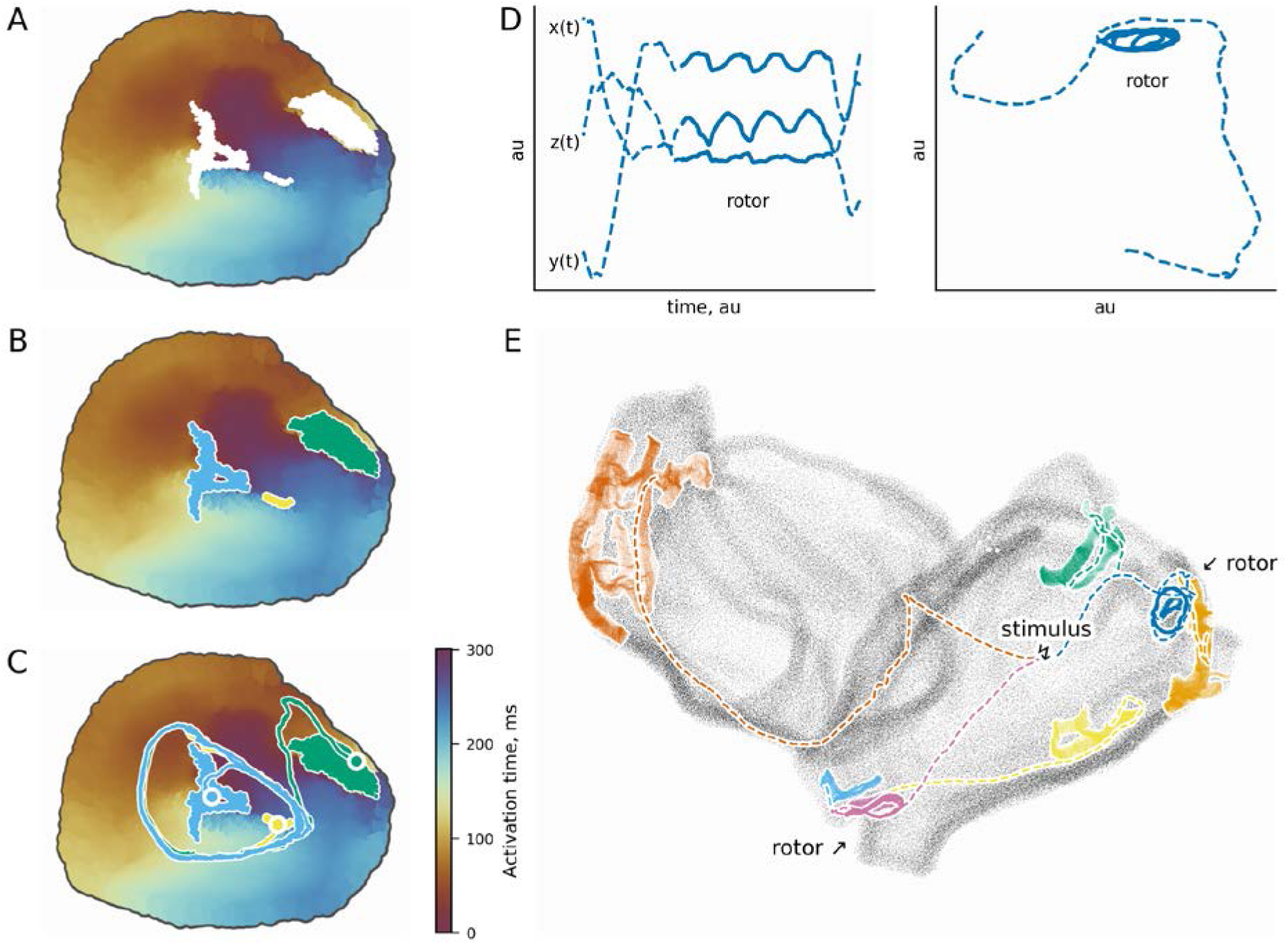
Backtracking algorithm pipeline of the reentrant driver search in 3D simulations. Wavebreaks (A), connected components (B), starting vertices and trajectories (C) for the small tissue region. Trajectory segmentation (D). Full pipeline (E). See text for details.

A single CPU comparison of computational time demonstrated that the RD algorithm is at least 700 times faster than the PS location via Hilbert transform. 2D simulations of 500 x 500 mesh demonstrated that phase map calculation via HT required more than 11.4±0.1 s, while consecutive PS location took 12.4±1.0 s. In contrast, the entire RD algorithm calculations took only 31.6±5.0 ms.

### Whole atria simulations

In total we have performed 320 whole atria simulations (2 heart geometries; 4 AP models; 40 pacing sites). As in previous computational and experimental studies (*16, 51*), we observed complex propagation patterns with colliding waves and numerous functional blocks that resulted in the generation of both stable and transient RDs (**Figure S6**). Due to the fact that both patient-specific geometries we used corresponded to persistent AF patients, AF was inducible in all AP models, including the SR model (**Figure 4**). We did not observe any notable differences in the frequency of AF occurrence between four cell-specific models, i.e. the number of pacing sites from which AF could be induced was similar, the difference was less than 10 % (**Table 1**). We have observed reentrant activity in 140 out of 320 simulations. In 14% of simulations, two RDs occurred simultaneously, in 9% – three or more. Some of the rotors were unstable (27%), they emerged or vanished in the middle of the simulation. Number of the RDs was the largest for the poAF (67) while other AP Models it was similar (**Table 2**).

**Table 1.** The number of pacing sites from which AF could be induced.

|  | persAF | pAF | poAF | SR |
| --- | --- | --- | --- | --- |
| Heart 1 | 9 | 9 | 12 | 14 |
| Heart 2 | 28 | 23 | 25 | 20 |
| Both | 37 | 32 | 37 | 34 |

**Table 2.**
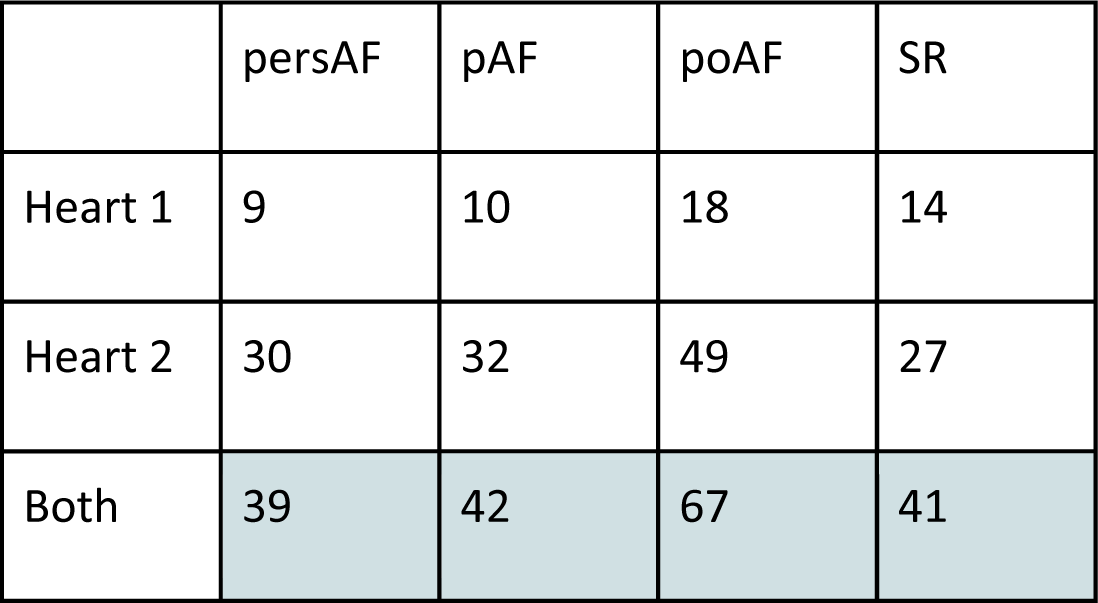
The number of rotors in 3D simulations.

|  | persAF | pAF | poAF | SR |
| --- | --- | --- | --- | --- |
| Heart 1 | 9 | 10 | 18 | 14 |
| Heart 2 | 30 | 32 | 49 | 27 |
| Both | 39 | 42 | 67 | 41 |

On the other hand, cell-specific models demonstrated notable differences in the prominence of wavebreaks. As demonstrated by shaded areas in **Figure 4A-B** the wavebreaks were mostly co-localized with the areas of dense fibrotic tissue (see also **Figure 5C** below). Thus, every AP model demonstrated functional blocks in the vicinity of pulmonary veins (both Heart 1 and Heart 2) and the right atrial appendage (Heart 2). However, in case of the SR model atrial regions susceptible to wavebreaks were confined to a small portion of the tissue (2.2% of the volume, **Table 3**); it was more extensive in case of other groups, persAF 17.5%; pAF 14.1%; poAF 18.7%. Moreover, in the case of persAF, AP model extensive fibrotic slabs were inexcitable in our simulations. These include portions of pulmonary veins (Heart 1, Heart 2) and RAA (Heart 2) (**Figure 4A**).

**Table 3.** Relative volume of the tissue susceptible to wavebreaks.

|  | persAF | pAF | poAF | SR |
| --- | --- | --- | --- | --- |
| Heart 1 | 6.2 | 5.3 | 7.2 | 0.8 |
| Heart 2 | 11.3 | 8.9 | 11.5 | 1.4 |
| Both | 17.5 | 14.1 | 18.7 | 2.2 |

As a consequence of these different effects of fibrotic substrate on the propagation patterns, we observed that RDs location was also AP-model dependent. PVs were arrhythmogenic in every AP model/geometry we used: in the case of the SR model RDs location was limited to pulmonary veins sleeves, while in the case of the pathological AP model arrhythmogenic region was extended to the myocardial tissue surrounding the veins (**Figure 4A**). The extensive fibrotic slab in RAA (Heart 2) was arrhythmogenic in SR, pAF, and poAF models. However, we did not observe any RDs there in the case of persAF model due to the partial inexcitability of this region. In the case of pathological AP models, we also observed a number (**Table 4**) of RDs close to the interatrial septum and mid-lateral RA.

**Table 4.**
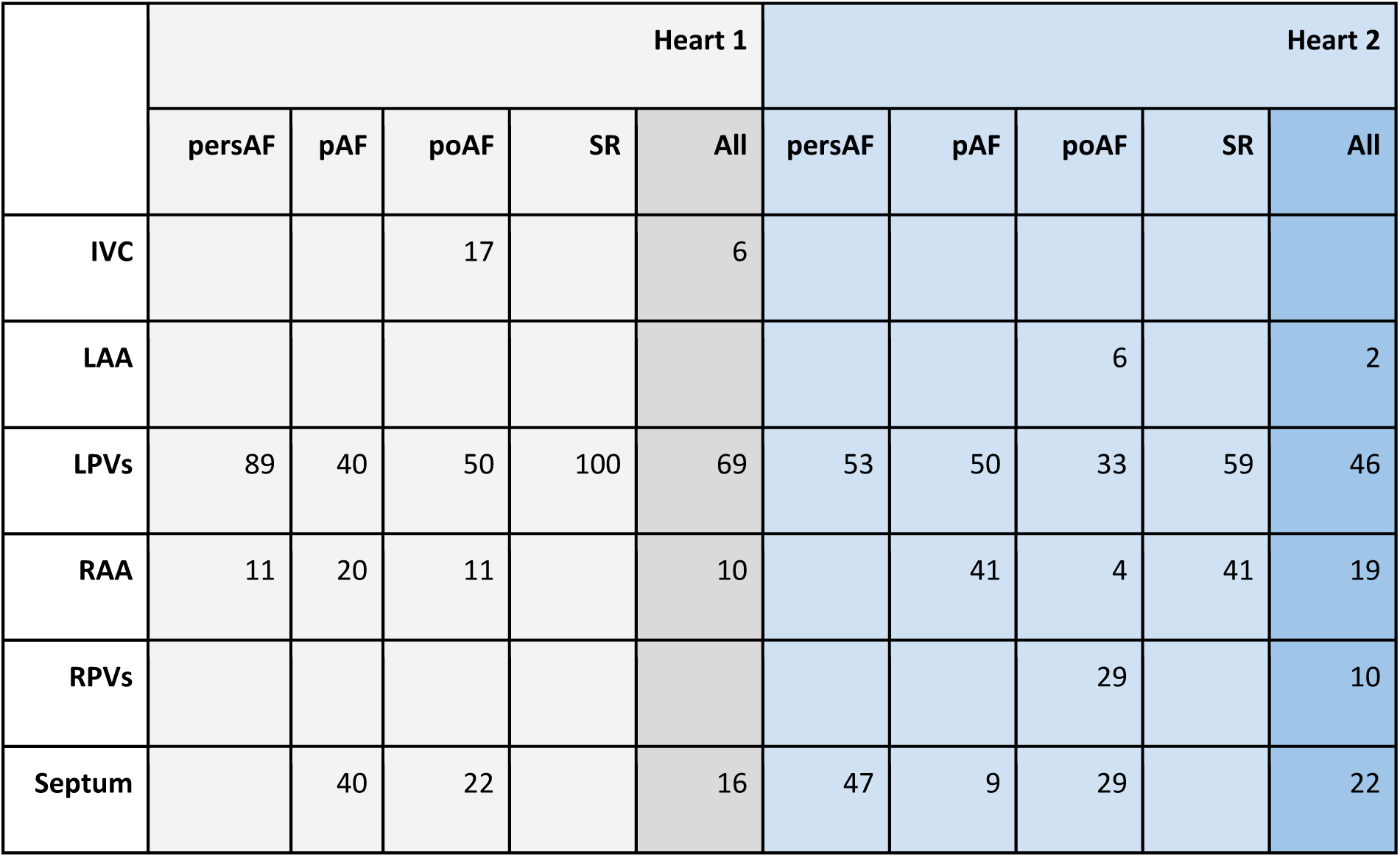
Rotor anatomical location. Value is a number of rotors in a particular location. %

**Figure 4C** demonstrates the RDs lifetime. The majority of the rotors were stable in the case of persAF model with a lifetime of 4 to 5 s. On the other hand, 27% of the rotors terminated in the case of the SR model before the end of the simulation. In the case of pAF and poAF models, 60 and 47% RDs correspondingly had a lifetime below 4 s, while some of these terminated (crosses in Fig4C), the others (circles in Fig4C) shifted from a different location. These results are in agreement with the transient nature of pAF. Other RDs characteristics, namely: rotation period, wavelength and conduction velocity are compared in Figure S7.

Thus, we can conclude that although RDs formed in a limited number of fibrotic regions, both the location of arrhythmogenic regions and their dynamics was AP-model dependent. In order to understand these differences, we have compared average wavebreak probability (**Figure 5A**) and fibrotic density (**Figure 5B**) within the scroll waves. More precisely, the former is the portion of tissue that is likely to produce a functional block; the latter is the ratio between fibrotic/nonfibrotic tissue. In the case of the SR model, the scroll waves rotate only around the dense fibrotic regions: from 39 to 94% of the tissue within the rotor was fibrotic. In the poAF model only, 18% of RDs rotated around areas with fibrosis density of less than 10%. The other two groups produced rotors in areas with fibrosis density from 12 to 50% (persAF) and from 10 to 88% (pAF). On the other hand, wavebreak probability was the lowest within arrhythmogenic regions for the SR model: persAF 0.026 ± 0.012; pAF 0.010 ± 0.004; poAF 0.010 ± 0.006; SR 0.004 ± 0.002 (**Figure 5A**). As shown in **Figure 5C,D**, there is a clear correlation between fibrotic density (FD) and wavebreak probability, however, the regression slope is different for different models: it is the lowest for SR model and the highest for persAF model. It should be noted that P(wb) was calculated over the first 2.5 seconds of the simulations, i.e. before the AF.

Due to the complex organization of atrial tissue, arrhythmogenic regions were not only model-dependent but also geometry-dependent, to take one example in the case of pAF FD was 0.27 ± 0.06 and 0.427 ± 0.0.20 for Heart 1 and Heart 2, respectively. On the other hand, distributions of P(wb) are similar for both patients as shown in **Figure 5A,E**: for example, in the case of pAF it was 0.008±0.004 for Heart 1 and 0.010 ± 0.004 for Heart 2. **Figure 5E** compares distributions of mean FD and P(wb) within 1-cm spheres in non-arrhythmogenic (gray circles) and arrhythmogenic regions (colored). One can see that RDs tend to reside within regions with higher P(wb). Interestingly, in the case of persAF, some regions with high FDs were not arrhythmogenic due to reduced excitability (Heart 2 in **Figure 5D**). This implies that while FD itself is an insufficient predictor of arrhythmogenesis, high P(wb) is a better metric that accounts both for structural and cellular remodeling. It should also be noted here that potentially arrhythmogenic regions could be estimated from a single simulation using the P(wb) metric. While 40 simulations with different pacing sites were required to directly locate RDs, P(wb) was mostly independent of the pacing site. See Figure S8 comparing P(wb) when atria were paced from the RAA and the LAA.

## Discussion

In this study we present whole-atria AF simulations incorporating personalized measurements of both structural remodeling from clinical imaging and electrophysiological remodeling from characterization of excised myocytes. Below we discuss in detail the two most important results of the article: firstly, the novel robust and computationally efficient algorithm of detecting the location of AF reentrant drivers (RDs), and, secondly, the clinical implications of simulation results.

### СV backtracking algorithm

In total, we have performed 320 whole atria computational EP studies in this investigation. Given the large number of simulations, manually tracking the RDs was tedious and unreliable. One of the most commonly used techniques in both computational and experimental studies is phase analysis (*22, 52*): the signal is transformed to the phase domain via Hilbert Transform and after that, RDs are tracked as a PS trajectory. However, this technique is computationally expensive and may result in both false positive and false negative RDs, when, for example, an excitation wave propagates along the scar or rotates around an anatomical obstacle (*23*). In order to overcome these limitations, we have developed a novel technique that consists of the following steps: (1) wavebreaks are localized, (2) CV field is backtracked from the WBs to the source of excitation, (3) quasi-periodic sources of excitation are identified by a neural network. The technique proves to be both robust (**Figure S3**) and computationally efficient. We compare the main advantages of the proposed technique below.

Firstly, while phase mapping requires voltage recordings, the BTA uses only the activation sequence as input data. This implies that this algorithm can be applied to various clinical and basic recordings performed on different platforms which provide activation sequence: optical mapping, monophasic action potential recordings, and extracellular electrograms (*53*).

Secondly, backtracking CV is computationally much more efficient, even on 2-dimensional 500×500 mesh it was 700 times faster than phase mapping. The difference is even larger in 3D due to the linear complexity of BTA. Indeed, to find the source of activation 1-dimensional subset of data is used, since the vertices of a single mesh element are being processed at each step. On the other hand, phase mapping requires the phase calculation for every vertex of the mesh first and then uses the whole phase map to locate phase singularities.

Thirdly, phase singularity is located in the electrotonically activated core of the reentry. As shown above, in **Figure 3**, myocyte oscillations on the PS trajectory are sub-threshold, while the actual source of arrhythmogenic activity is the attractor of CV trajectories. Consequently, phase mapping might underestimate the size of the arrhythmogenic region since the actual RDs cover a much larger area than the PS trajectory. Moreover, in some cases of drivers, such as reentry rotating around an anatomical obstacle or a focal source of activation, an arrhythmogenic source of activation can be located by the BTA, but not by the phase mapping technique.

**Figure 3.**
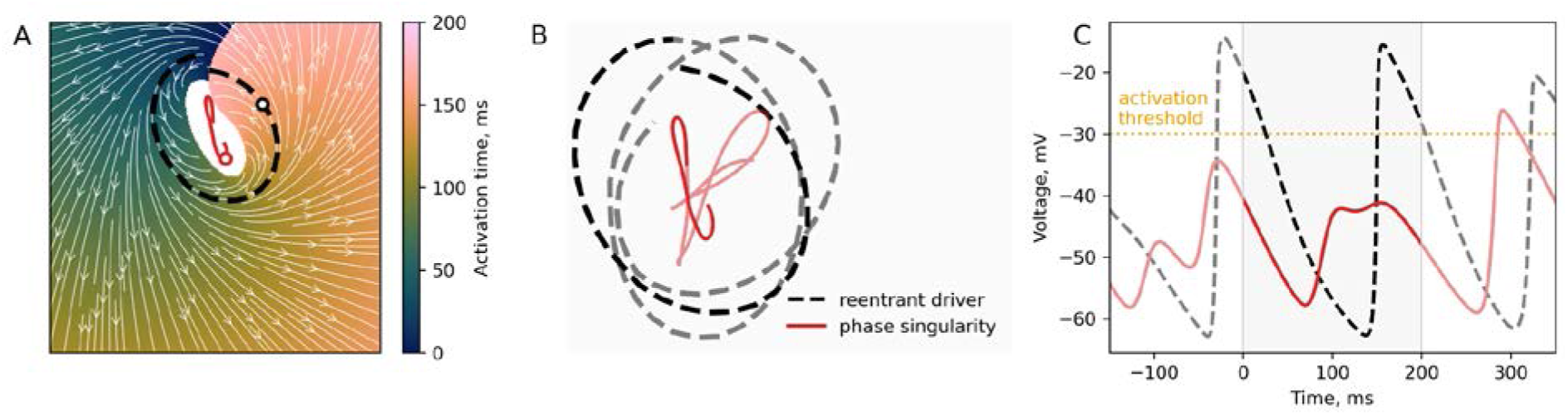
Comparison of the algorithms of the rotor trajectory search: 2D simulations. A. Activation sequence for single rotation period. The color represents the activation time, and the white arrows represent the conduction velocity field. We define the activation when transmembrane voltage exceeds the threshold of -30 mV. The white area is the inexcitable tissue. The dashed black line and the solid red are the trajectories obtained using BTA and the Hilbert transform, respectively. The latter produces the trajectory within the inexcitable core with subthreshold depolarizations of the transmembrane voltage. B. The same trajectories for several rotation periods. C. Transmembrane voltage for the points marked by circles in panel A. Gray background shading (0 -- 200ms) corresponds to the rotation represented in panel A. The phase singularity trajectory (red) is always below the activation threshold.

Previously, Vandersickel et al. proposed another technique named Directed Graph Mapping (DGM) which was successfully applied to both simulated (*25*) and clinical recordings (*26*) of reentrant activity. DGM builds a directed graph from the activation sequence and then searches for the shortest possible closed loop in the graph. The algorithm we propose here has several important advantages. First, DGM uses the breadth-first search (BFS) algorithm in order to find cycles, thus moving forward in time. The technique we propose here moves backward in time, from present to past, building a single trajectory from a given starting point. Moving backward in time is computationally more efficient and also more robust: as shown in **Figure 3A** minor deviations from the RD would result in the trajectory moving away from the RD towards the core or the outer part of the tissue. Secondly, DGM seeks for the shortest possible cycle (rotor) in the graph. Similarly to phase analysis, the DGM technique might thus underestimate the arrhythmogenic region, which might be important in the case of computationally guided ablation procedures. For example, a minimal reentrant loop would encircle the white area in **Figure 3A**.

### Patient specific simulations

The gradual progression of AF from paroxysmal, to persistent, and to permanent is currently understood via the “AF begets AF” hypothesis (*2, 3*). Specifically, persistent AF may result from remodeling on two levels: electrophysiological and structural remodeling (*44, 45*). The former increases the stability and frequency of rotors because of AP shortening (*2, 8, 46, 48*), hyperpolarization of the resting membrane potential (*54, 55*), and connexin downregulation (*56*). The latter contributes to alterations in CV and the shape of the wavefront, forming rotors, in addition to fibroblast proliferation and collagen deposition (*57*– *59*).

The main aim of our study was to investigate the role of electrophysiological remodeling in AF. To keep all other factors equal, we have combined the same two patient-specific persAF geometries with four different pathology-specific AP models. Although this combination is strictly theoretical, it provides valuable insights, because it is almost impossible to separate different kinds of remodeling in actual human heart experiments.

One classical pharmacological approach to treat AF is to reverse electrophysiological remodeling, for example, class IC antiarrhythmic flecainide that both prolongs AP and partially blocks ryanodine receptors (*60*). More recently, targeting atrium-specific I_K,ACh_ was proposed (*61–63*), however despite promising animal model experiments, human studies were disappointing (*64*). Our results concur with limited efficacy of pharmacological ‘normalization’ of AP in AF. On one hand, in our simulations the break-up of the wavefront is the most frequent in the case of persAF AP model (**Figure 5**). The tissue that is prone to wavebreak is also most extensive in this case (**Figure 4**). In the case of the SR AP model the probability of wavebreak was the lowest and limited to few fibrosis-heavy regions, implying reduced generation of spiral waves in the last case. However, despite apparent difference in the frequency of wavebreaks we did not observe any difference in the frequency of arrhythmogenic RDs generation between the four pathology-specific models. Our results thus indicate that in case of heavy fibrosis reversing electrophysiological remodeling would not revert AF inducibility. However, as we will discuss below, there is a pronounced difference in the location of arrhythmogenic regions.

**Figure 4.**
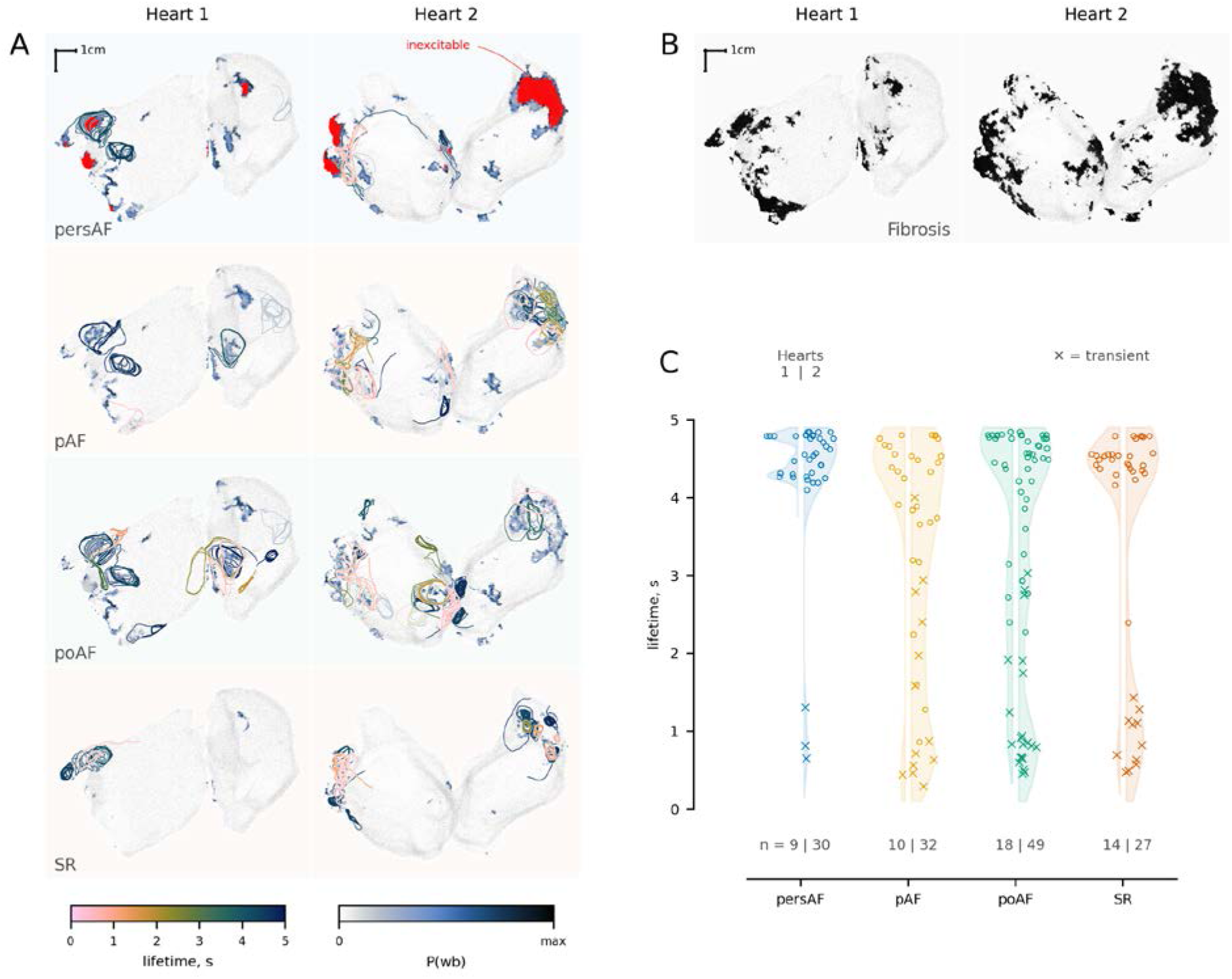
Reentrant drivers in 3D. A. Four rows correspond to four AP models. The left and right columns correspond to Heart 1 and 2, respectively. Rotor trajectories are superimposed over the geometry and colored according to their lifetime. Transparent rotors are on the back side of the heart. Blue-shaded regions show the vertex-wise wavebreak probability, P(wb). Inexcitable areas are colored in red. B. The spatial fibrosis distribution. C. The lifetime of the reentrant drivers. The Left and right halves of the violins are Heart 1 and 2 correspondingly. “x” symbol signifies transient rotors.

Haissaguerre et.al. (*10–12*) were the first to report the role of pulmonary veins in pAF. Despite the success of PV isolation in pAF it was less satisfactory in the case of persistent AF (*65, 66*) due to wider atrial substrate. In a later study, Haissaguerre et. al. reports the distribution of RDs in the study of 103 patients with persistent AF (*13*). The RDs were mainly located around left and right PVs: if 21 patients with prior PV isolation were excluded, the prevalence of RDs in these regions accounted for 94 and 97%, correspondingly. However, apart from PV regions, 81% of patients also had RDs in the inferior posterior left atrium, approximately half of the patients had RDs in the RAA or anterior left atrium, ∼10% had RDs in the IVC or interatrial septum. The distribution of RDs in our simulation corresponds to a wide atrial substrate in persistent AF (**Table 4**). PVs were arrhythmogenic in both geometries and for every single AP model, accounting for 46% of RDs. However, RAA and septum were also arrhythmogenic, respectively, accounting for 14% and 25 % of RDs.

Although the frequency of arrhythmias was not model-dependent, the distribution of RDs depends heavily on the AP model. SR model demonstrated a limited number of arrhythmogenic regions: it was either PV sleeves or, in the case of Heart2 geometry, fibrotic-heavy RAA. In the case of pathological AP models, the arrhythmogenic region extended to the tissue surrounding PVs and interatrial septum. Surprisingly, we observed the most extensive arrhythmogenic region not in the case of persAF AP model but in the cases of pAF and poAF models (Figure 4). That may be explained by the reduced excitability of persAF model: indeed unidirectional functional block is required in order to initiate the reentrant wave (*44, 67, 68*); however, in the case of persAF, we observed several regions that were completely inexcitable, essentially resulting in bi-directional block.

Despite somewhat less extensive arrhythmogenic region in case of the persAF, as expected (*2, 8, 46, 48*), shorter APD did stabilize the scroll waves: vast majority of RDs were stable in the case of persAF model, however 25-60% of RDs were transient in other cases (Fig.4C). Moreover, in our simulations more extensive atrial substrate resulted in the emergence of several simultaneous RDs: in the cases of SR and persAF *85-95%* of arrhythmias were driven by a single RD, while in the cases of pAF and poAF *29* and *46 %* of arrhythmias correspondingly were driven by two or more RDs.

To explain these pathology-specific differences, we have calculated the wavebreak probability (Figure 5). Previously (*16*) Shannon’s Entropy was proposed to predict pro-arrhythmogenic regions. It was shown that RD tends to reside not in the densest fibrotic areas, but in the border zone between fibrotic and healthy tissue, where both fibrosis density and fibrosis entropy was high. This metric, however, does not account for electrophysiological remodeling and thus cannot explain differences between pathology-specific models that we observe in our study. In this article we demonstrate (Figure 5) that arrhythmogenic areas tend to colocalize with regions of high P(wb). It is important to note, that since we were interested in a predictor of arrhythmogenesis, the wavebreak probability was calculated using the initial part of simulations, before the generation of RDs. Indeed, wavebreak was defined as a delay in wavefront propagation (more precisely as CV < 1 cm/s). In these terms, scroll waves essentially rotate around the wavebreak region, thus including the corresponding part of simulations would distort distribution in Figure 5E.

**Figure 5.**
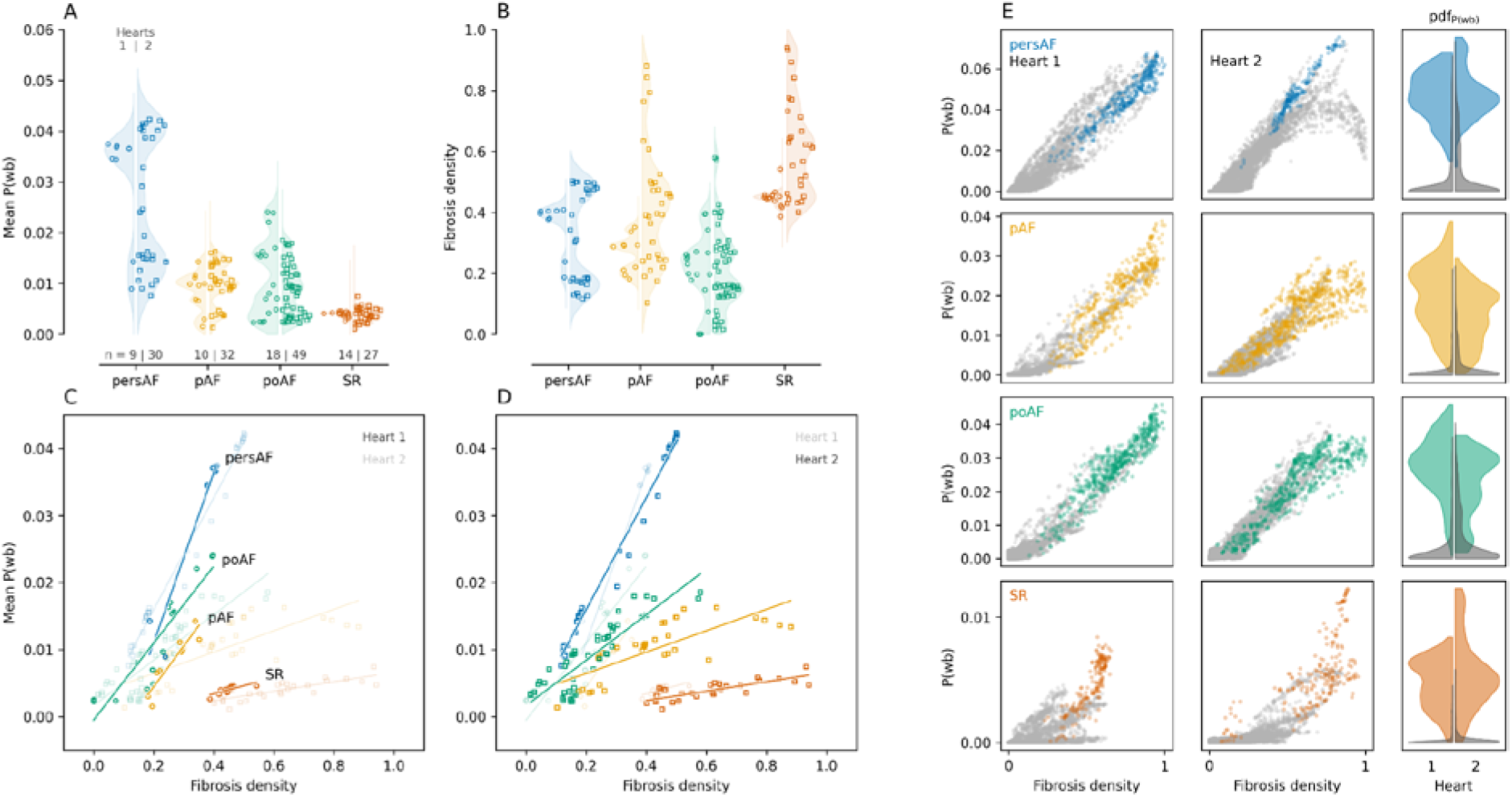
Mean P(wb) and fibrosis density in 3D simulations. A-B. Mean wavebreak probability, P(wb), (A), and fibrosis density (B) encircled by scroll waves. Each dot corresponds to a different reentrant driver. More precisely, the values were calculated within (1) tissue within a 0.5 cm radius from the reentrant driver trajectory and (2) tissue surrounded by the rotor. C-D. Correlation between mean P(wb) and mean fibrosis metrics for the Heart 1 and Heart 2 geometries. E. Each color-coded dot in swarm plots corresponds to a 1 cm-sphere within the arrhythmogenic regions (inside scroll waves); gray dots correspond to the remaining (non-arrhythmogenic) tissue. Corresponding probability distributions of P(wb) are shown on the rightmost panels.

Our results indicate (Figure 5C, D) that there is a clear correlation between the fibrosis density and P(wb); severe electrophysiological remodeling increases the regression slope, which explains more extensive arrhythmogenic substrate in the pathological models. On the other hand, in the case of the persAF model, extreme fibrotic density reduced the activation probability and, consequently, the wavebreak probability (Figure 5E, top-right panel), which explains the relatively modest arrhythmogenic substrate in the case of persAF model in comparison with pAF and poAF models.

In our study, the fibrotic tissue was simulated by reduction of I_Na_, I_CaL_, I_K1_ and gap junction conductivities (*16, 69, 70*). Previous simulations (*68, 71*) in which fibrotic tissue was a non-conductive obstacle demonstrated similar results. On the one hand, increasing fibrosis density above a certain threshold reduces percolation probability and, consequently, the RD probability. On the other hand, this threshold is affected by reduced excitation, shifting arrhythmogenic regions away from dense fibrotic regions. Arguably, this indicates that, for example, sodium channel block by class 1C agents (*72*) could contribute to AF treatment despite slowing of conduction: at least in some cases, reduced excitability might also reduce the potentially arrhythmogenic region.

### Study limitations

Patient-specific models were reconstructed from the AP recorded in myocytes isolated from RAA. This implies that we did not account for region-specific phenotypic differences in atria. Although these differences are important and affect arrhythmogenic substrate, it should also be noted that left-to-right gradient is reduced in persAF (*73*). The spontaneous calcium releases resulting in delayed afterdepolarization are shown to occur more often in AF (*74*). These may contribute to wavebreaks but were not considered in this study.

### Conclusions

In this study we demonstrate whole-atria simulations in which patient-specific models were used to account for both structural and electro-physiological cellular remodeling. To reconstruct heart geometry, both the fiber orientation and the areas of fibrosis, DGE-MRI data from two persAF patients were used. To reconstruct pathology-specific AP models, we used myocytes excised from the RAA of four different patients. We were blinded to the patient’s diagnosis, only the AP measurements were provided to us. Although we used an artificial combination of persAF tissue remodeling with pathology-specific electrophysiological remodeling, our article provides the pipeline for patient-specific simulation of AF and answers several questions. While previous computer simulation studies successfully predicted the location of arrhythmogenic regions in atria (*15*), the EP remodeling was not personalized, and it was unclear to what degree RD locations depended on AP morphology and restitution. We observed the largest number of wavebreaks in the case of persAF AP model and the lowest in the case of SR model. Four patient-specific models had common arrhythmogenic regions (PVs and RAA), but the AP models corresponding to persAF, pAF and poAF had more distributed arrhythmogenic substrate. However, the arrhythmogenic region was most extensive in pAF and poAF simulations due to the reduced excitability of persAF AP model. We also demonstrate that RDs tend to colocalize with the highest wavebreak probability regions. The wavebreak probability correlates with fibrosis density, but these regions also depend on electrophysiological remodeling. Lastly, in order to analyze the RDs location and lifetime, we developed a novel algorithm based on backtracking the CVs vector field, thus requiring only the activation sequence as input data, which we demonstrate to be computationally efficient and robust. The advantages of the algorithm imply that it can be used for experimental and clinical recordings as well; we plan to address these possible applications in our future studies.

## Materials and methods

### Recordings of cellular action potentials in AF patients

Right atrial appendages were obtained from patients undergoing cardiac surgery. Experimental protocols were approved by the ethics committee of the University Medical Center Göttingen (No. 4/11/18), and informed consent was obtained from each patient. The patient samples were grouped in four blinded sets according to the diagnosis: Sinus Rhythm (SR), postoperstive AF (poAF), paroxysmal AF (pAF), and persistent AF (persAF). Episodes were classified as pAF if they terminate spontaneously within seven days, or persAF, if the episode lasted longer than one week (*27*). Sinus rhythm patients were followed-up for 6 days to identify those with postoperative AF (poAF, any episode of AF lasting longer than 30 s) and without postoperative AF, classified as SR. Excised right atrial appendages were subjected to a standard protocol for myocyte isolation (*6*). Right atrial myocytes were suspended in EGTA-free storage solution for subsequent simultaneous AP measurements.

Only rod-shaped myocytes with clear striations and defined margins were investigated. During experiments, myocytes were perfused with a bath solution at 37 °C containing (in mmol/L): CaCl_2_ 2, glucose 10, HEPES 10, KCl 4, MgCl_2_ 1, NaCl 140, probenecid 2; pH=7.35. The pipette solution contained (in mmol/L): EGTA 0.02, GTP-Tris 0.1, HEPES 10, K-aspartate 92, KCl 48, Mg-ATP 1, Na_2_-ATP 4; pH=7.2. To evoke APs, 1 ms current pulses of 1.5–2 threshold strength were applied. The Pacing Cycle Length (PCL) was decreased in a stepwise manner, starting at 2064 ms and subsequently 1032, 516, 344, 258, 206, 172, 148, 129 ms. Pipette resistances in the range of 5–7 MΩ were utilized.

Due to computational limitations, a single patient from each group was chosen arbitrarily for model optimization (see below “Genetic algorithm”). Patients’ data is given in **Table S1**, and APs restitution is shown in Figure 1B. All patients were males undergoing coronary artery bypass surgery.

### Acquisition of atrial anatomy in AF patients

Cardiac magnetic resonance imaging (MRI) was performed on a 1.5T scanner (Magnetom Avanto, Siemens Medical Systems, Erlangen, Germany) equipped with a 32-channel cardiac coil. LGE-MRI was performed 15 min after administering gadolinium chelates using a 3D, ECG-gated, respiratory-navigated, and inversion recovery-prepared Turbo Fast Low Angle Shot sequence with fat saturation (voxel size: 1.25 × 1.25 × 2.5 mm^3^). In the resulting images, the biatrial wall was manually contoured, and LGE and non-LGE regions were segmented using an adaptive histogram thresholding algorithm as described previously (*16*). The fibrosis burden derived from LGE segmentation was expressed as a percentage of the atrial wall. Segmented images were up-sampled to an isotropic voxel size of 400 mm^3^ using shape-based interpolation, and 3D finite element meshes were generated from the resulting high-resolution datasets using a previously developed approach. In each patient-derived model, myocardial fiber orientations were assigned using a rule-based method (*28*).

At the tissue level, conductivity values were assigned so that an effective longitudinal conduction velocity of 43.39 cm/s was achieved in the non-fibrotic myocardium, which was within the range of values recorded in patients with AF (*29*). Fibrotic regions were represented with remodeled electrophysiology, anisotropy, and conduction properties. The non-fibrotic action potential model was modified as described previously (*16*) to represent the regional electrophysiological changes due to fibrotic remodeling: 50% reduction in inward rectifier potassium current I_K1_; 50% reduction in L-type calcium current I_CaL_; and 40% reduction in sodium current I_Na_. Conductivity values in fibrotic regions were reduced by 30% to represent decreased intercellular coupling due to replacement fibrosis, collagen deposition (interstitial fibrosis), and gap junction remodeling. Since fibrosis results in greater conduction velocity impairment in the direction transverse to cardiac fibers, the conductivity values were further modified to achieve a longitudinal to transverse anisotropy ratio of 8:1 (*15*).

Electrical wave propagation was governed by the monodomain formulation, and finite-element simulations were executed with the CARP software package(*30*); numerical detail can be found in previous publications (*31, 32*). In each patient-derived model, 40 pacing sites were distributed uniformly throughout the atria. At each pacing site, a clinically relevant programmed electrical stimulation pacing sequence 34 of 14 stimulation pulses with cycle lengths decreasing from 300 to 150 ms in 25 ms intervals was applied to induce AF and assess the arrhythmogenic propensity of the fibrotic substrate. For each of the patient-derived models, we simulated 40 AF induction protocols, 1 for each pacing site (*15, 16*).

### Genetic Algorithm

Maleckar AP model (*33*) was used in the study with the following modifications. Ventricular ICaL current (I_CaLv_) was added to the original Maleckar model as described in our preliminary study (*34*) in order to replicate phenotypic variability among AF patients as observed in experimental AP recordings. The expression of the ventricular isoform of L-type calcium channel in AF is supported by RNA-seq studies (*35*) and our recent CAGE study (*36*). I_CaLv_ was simulated as the modification of the I_CaL_ current in the O’Hara-Rudy model (*37*) modified by Tomek (*38*). AP model is integrated using LSODA solver (*39*). Relative and absolute tolerances were set as 10^-4^ and 10^-7^, respectively.

AP model parameters were personalized to conform to the patch-clamp recordings *via* custom Genetic Algorithm modification (*40*). Briefly, an initial generation of the organisms (different model parameter sets) is iteratively updated via selection, mutation, and crossover genetic operators to minimize the deviation between simulated and experimental AP waveform restitution. As described previously, only steady-state AP waveforms were considered the “proper” solution to the optimization problem. This was achieved by the inclusion of intracellular concentrations in the set of optimized parameters. The full list of the optimized parameters with their boundaries is provided in **Supplementary Text**. A logarithmic scale was used for ionic channel conductivities during the optimization process. Optimized model parameters and AP markers for the model and experiment are in **Table S2**.

### Backtracking algorithm

We have developed a novel algorithm to locate the reentrant drivers. The algorithm consists of the following stages (**Figure 2**).

1. First, we calculate the distance (*h*) and activation time delay (Δ*t*) between neighboring vertices. For every activated vertex, we define the wavebreak as the event when *h/*Δ*t < 1 cm/s* between any neighboring vertices. Such wavebreaks are depicted as white dots in **Figure 2A**. In all our simulations CV was above 5 cm/s.
2. Two wavebreaks are ascribed to the same connected component (in 4D = 3D + time) if these were close in space and happened almost simultaneously, more precisely: *h_wb_/*Δ*t_wb_ > 1 cm/s*. Three connected components are exemplified in **Figure 2B** by different colors.
3. For every component, we find the last activated vertex (colored circles in **Figure 2C**). The algorithm proceeds by building a trajectory by backtracking the velocity field (colored curves in **Figure 2C**). Since the core of the vortex (*41*) is identified as a wavebreak on step 1, every reentrant driver can be located as a part of the trajectory starting from the corresponding core (**Figure 3A**). The conduction velocity was calculated as the inverse gradient of activation time: 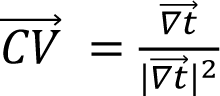. Detailed algorithm of CV calculation is provided in the **Supplementary Text.**
4. The trajectories from step 3 should be segmented in two parts: quasiperiodic rotating source of excitation (solid colored line in **Figure 2D**), and the ‘tail’ connecting it with the wavebreak component. This is done via the neural network trained on synthetic trajectories (see **Supplementary Text and Figure S1**). We have used a modified U-Net to solve the segmentation problem (*42*). Modifications to the U-net are described in the **Supplementary Text,** neural network architecture is shown in **Fig S2.** The straightforward approach to the problem is the direct segmentation of trajectory, *i.e.* using 𝑟𝑟⃗ (𝑡𝑡) as input to the neural network. However, in this case, even a relatively small number of misclassified points would result in a large number of artifact rotors. Instead of that we used the matrix of pairwise euclidean distances between the trajectory points was used as input to neural network, *i.e.* 𝐴𝐴(𝑡𝑡_1_, 𝑡𝑡_2_) = || 𝑟𝑟⃗ (𝑡𝑡_1_) − 𝑟𝑟⃗ (𝑡𝑡_2_)||_𝐿𝐿2_ (**Figure S1C**). The output matrix elements were supposed to be ***1*** if both 𝑟𝑟(𝑡𝑡_1_) and 𝑟𝑟(𝑡𝑡_2_) belong to the rotor segment and ***0*** otherwise. Finally, each minimal diagonal block of the output matrix was interpreted as corresponding to a *single* reentrant driver if it contained a *single* connected component of non-zero elements. This 2D matrix representation is very robust: as shown in **Figure S3** even **35** % of misclassified matrix elements result in correct trajectory segmentation. Segmentation of synthetic datasets (**Table S3**) resulted in 95% accuracy and 93% IoU (intersection over union). Segmentation of an actual BTA trajectory recovered from whole atria simulations and its 2D matrix representation is exemplified in Figure S4.

In order to compare new BTA with the traditional Hilbert Transform PS algorithm (*43*) we have simulated the reentry in 2D square tissue, 5 x 5 cm. Simulation lasted 1 s. The tissue was discretized in a rectangular grid with a 100 um step that resulted in 500 x 500 nodes. Transmembrane voltage was recorded at every 1 ms. SR AP model was used for all nodes. Both algorithms were implemented in the Julia programming language without multiprocessing.

#### Wavebreak probability, *P(wb)*

For every vertex in the mesh, we calculate wavebreak probability as a proportion of the stimuli resulting in wavebreaks (defined as above: CV below 1 cm/s). Only the first 2.5 s of the simulations were considered for the probability estimate, that is, when the tissue was stimulated by a virtual pacing electrode and not by reentrant activity. We have to note here that *P(wb)* defined as above is low in the regions with reduced excitability, more formally: *P(wb)=P(wb*|*act)*P(act)*, where *P(act)* is the activation (depolarization) probability of the vertex, *P(wb*|*act)* is the conditional wavebreak probability, provided the vertex was activated (**Figure S5**).

In order to account for non-local properties of the fibrotic tissue, we calculated mean 𝑃𝑃(𝑤𝑤𝑤𝑤) = Σ(<u>^𝑣𝑣^</u> ⋅ <u>^𝑁𝑁𝑤𝑤𝑤𝑤^</u>) as a weighted sum of the fractions, <u>^𝑁𝑁𝑤𝑤𝑤𝑤^</u>, over the tetrahedrons constituting the volume. 𝑁𝑁_𝑤𝑤𝑤𝑤_ is the total number of the wavebreaks, 𝑁𝑁_𝑠𝑠𝛻𝛻𝑠𝑠𝑠𝑠_ is the total number of stimuli (480). The latter equals the number of simulations (40) times the number of stimuli per simulation (12). Weight is the tetrahedron volume fraction,

## Abbreviations

AF: atrial fibrillation
AP: action potential
persAF: persistent AF
pAF: paroxysmal AF
poAF: post-operative AF
CV: conduction velocity
FD: fibrotic density
PS: phase singularity
PV: pulmonary vein
RA: right atrium
RD: reentrant driver
SR: sinus rhythm
RAA: right atrial appendage
LAA: left atrial appendage
BTA: backtracking algorithm

## Acknowledgments

**Funding:** Leducq Foundation grant RHYTHM to IE and NT, National Institutes of Health grant R01-HL141470 to IE, National Institutes of Health grant R01HL166759 to NT, Deutsche Forschungsgemeinschaft (DFG, VO 1568/3-1, VO1568/4-1, SFB1002 project A13 and under Germany’s Excellence Strategy—EXC 2067/1—390729940) to NV, German Center for Cardiovascular Research (DZHK 81X4300102, 81X4300115, 81X4300112) to NV.

**Author contributions:** Conceptualization: RS, NV, NT, IE. Software: AVP, RS, RA, AP, PMB. Investigation: AVP, VS, RA, AP. Visualization: AVP. Supervision: RS, NV, NT, IE. Resources: IK, NV, NT. Writing—original draft: AVP, RS. Writing—review & editing: AVP, RS, RA, AP, PMB, VS, IK, ER, NV, NT, IE.

## Supplementary Materials for

**This PDF file includes:**

Supplementary Text

Tables S1 to S4

Figs. S1 to S8

## Supplementary Text

### Genetic algorithms

The parameter values were fitted via GA as in (40). In the current study, we used 1000 generations, 8192 organisms, and 128 elites. The following concentrations were set to the fixed experimental levels: [Mg2+]_i_ 1 mM, [Na+]_o_ 140 mM, [Ca2+]_o_ 2 mM, [K+]_o_ 4 mM. The following parameters were subjected to genetic operators: conductivities of INa, atrial and ventricular phenotypes of ICaL, Ito, IKur, IK1, IKr, IKs, IBNa, IBCa, INaK, ICaP, INCX, RyR, SERCA and [Na+]_i_, [K+]_i_, [Ca2+]_rel_ concentrations. Ionic current conductivity bounds were set from 0.1 to 10 times the original model value, except for INa (0.8, 10.0) and ICaL (0.01,10.0). [Na+]_i_ was mutated from 4 to 16 mM, [K+]_i_ from 100 to 160 mM, [Ca2+]_rel_ from 0.06 to 6.0 mM.

The following CLs were used: 2064, 1032, 516, 344 ms for persAF, poAF, SR; 516, 344, 256, 206 for the pAF. Longer periods for pAF trace were excluded due to experimental artifacts in the corresponding recordings.

The code of the GA can be found at https://github.com/humanphysiologylab/ga-meta.

### CV calculation

We calculate velocity, 𝐶𝑉, for the current tetrahedron given the activation times, 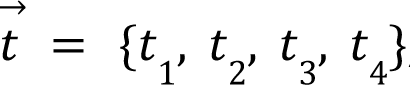, of the vertices and their coordinates, 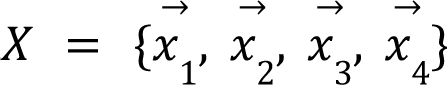.

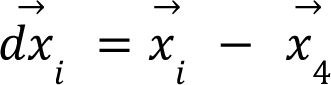

𝑈 – 3×3 matrix, corresponding to the projection operator on the tetrahedron edges:

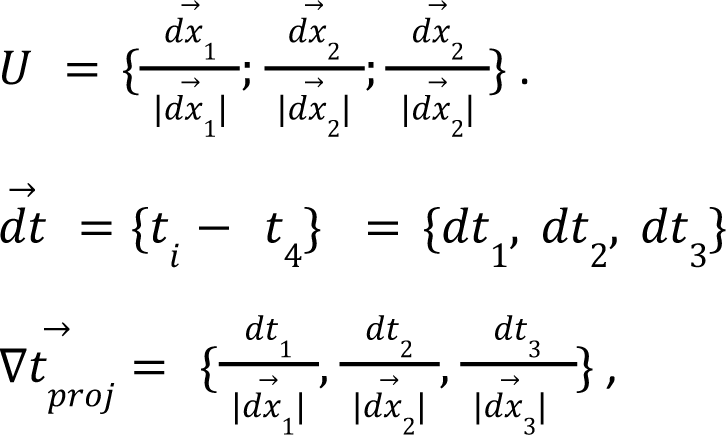

where 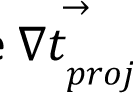 – projections of the 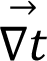 onto tetrahedron edges; 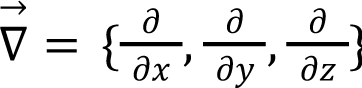.

Then activation time gradient 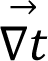 is the solution of 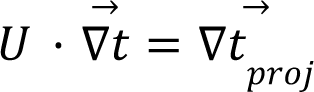.

The conduction velocity can then be calculated as the inverse gradient of activation time: 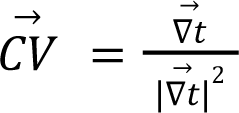

The code can be found at https://github.com/humanphysiologylab/heart-meshes/blob/1e94c4c71bdd7f2910d8798056e81426044251ac/src/mesh-free/calculate_cv.jl.

### Synthetic trajectories

Neural network was trained via a supervised learning algorithm. Since manual markup is tedious, we created a synthetic dataset that was marked up automatically (upon generation). Synthetic trajectories combined two regimes: rotor (quasi-periodic behavior) and non-rotor (non-periodic curve).

Rotor segments of trajectory were modeled as a modulated combination of harmonic signals:

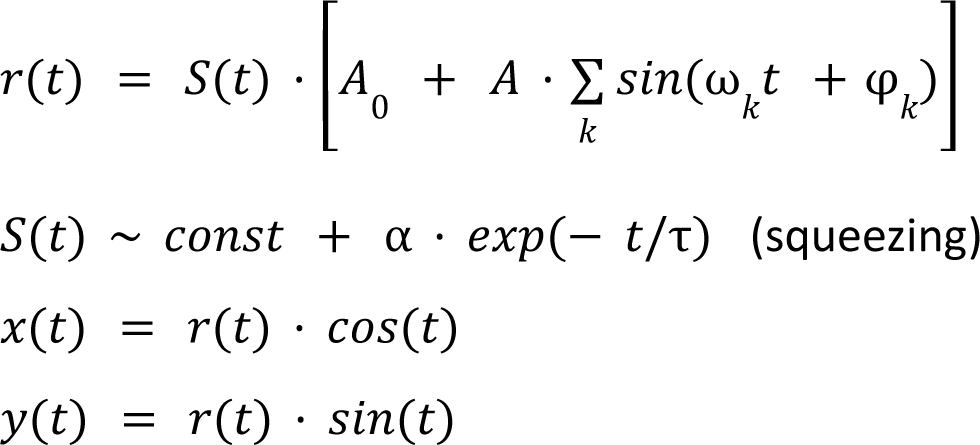

Non-rotor segments were modeled as a random walk with the following smoothing via the moving average

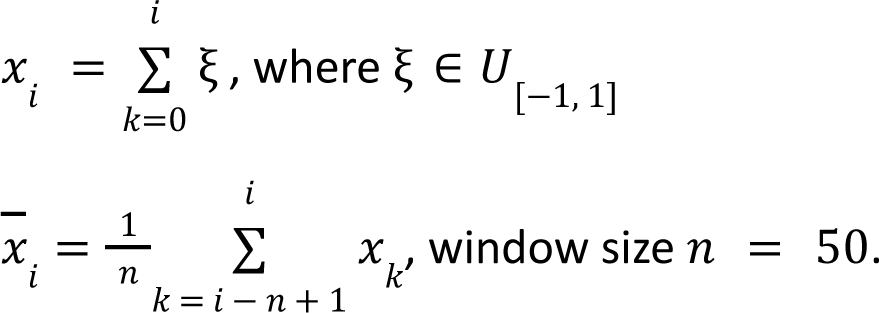

All parameters were set manually to produce trajectories similar to the experimental ones. To generate the final trajectory, we concatenated rotor and non-rotor segments as follows: rotor → non-rotor → rotor → non-rotor, etc. New segment was added to the trajectory after translation and rotation in order to make the connection smooth (**Figure S1 A,B**).

The code can be found at https://github.com/humanphysiologylab/nn-rotor/blob/master/notebooks/001-Synthetic-rotors.ipynb.

### Trajectory segmentation

The straightforward approach to the trajectory segmentation is using 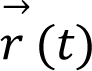 as input to the neural network. However, this results in a number of undesirable artifacts: for example, a single misclassified pixel might split the reentrant driver in two, and consequently, the number of artifact rotors is increased dramatically, while their lifetime is reduced. Instead of that we used the matrix of pairwise euclidean distances between the trajectory points was used as input to neural network, *i.e.* 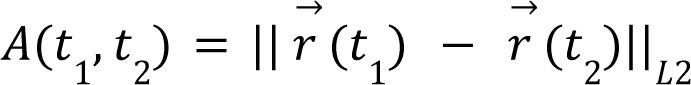 (**Figure S1 C,D**). The output matrix elements were supposed to be ***1*** if both 𝑟(𝑡_1_) and 𝑟(𝑡_2_) belong to the rotor segment and ***0*** otherwise.

The following algorithm was used for the trajectory segmentation using its 2D representation (Figure S3):

(1) Fix some time 𝑖 and duration 𝑛.

(2) Calculate the mean number of pixels classified as rotors, 𝑝, over the edge of the 𝑛 × 𝑛 square with a center at the diagonal (𝑖; 𝑖).

(3) If 𝑝 is greater than a threshold (we set 0.75) then mark trajectory segment from 𝑖 − 𝑛/2 to 𝑖 + 𝑛/2 as a rotor.

(4) Iterating over all possible 𝑖-s and 𝑛-s, we merge all rotor segments with nonzero intersection.

### Neural network architecture

We have adapted the U-net model (4*2*) (**Figure S2**). The code can be found at https://github.com/humanphysiologylab/nn-rotor.

The network consists of two mirrored cascades: the downsampler and the upsampler. The downsampler iteratively halves spatial resolution (via max-pooling layers) and doubles the number of channels. The upsampler does the opposite job, doubles spatial resolution (via upsampling layer) and halves the number of channels. Both parts share the same core block which is a twice-repeated sequence of the following layers: convolutional (3×3 kernel), batch-normalization, and activation (ReLU). There are residual connections between the downsampler and upsampler.

The network’s input is a gray-scale square image while the output is a matrix of the same size, matrix elements are probabilities of the target class in the same position in the input image.

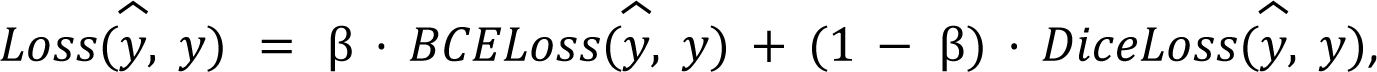

where 𝑦 and 𝑦 are predicted and true probabilities. Such loss function is also known as “ComboLoss”.

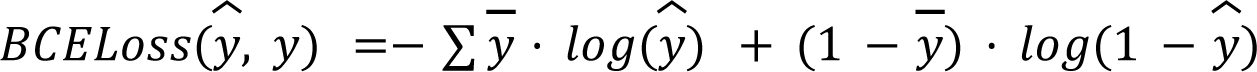

BCELoss is Binary Cross Entropy Loss (for numerical stability, we used BCE with logits). Label smoothing was used for the true labels in BCELoss:

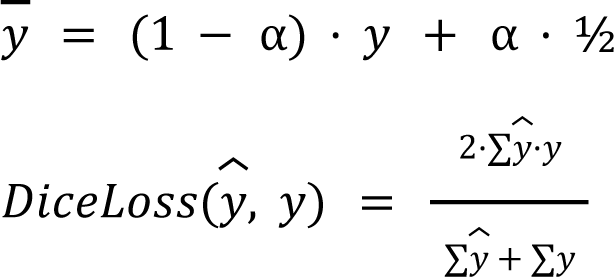

We used the following hyperparameters for loss function: α = 0. 1, β = 0. 5.

**Table S1.** Patients’ information.

|  | **SR** | **poAF** | **pAF** | **persAF** |
| --- | --- | --- | --- | --- |
| Sex | male | male | male | male |
| Age, y | 72 | 59 | 62 | 77 |
| Body mass index, kg/m<sup>2</sup> | 26 | 27 | 41 | 23 |
| CAD | Y | Y | Y | Y |
| MVD/AVD | N | N | N | N |
| Hypertension | Y | N | Y | Y |
| Diabetes | Y | N | N | N |
| Hyperlipidemia | N | N | Y | Y |
| LVEF,% | 64 | 32 | 35 | 45 |
| Digitalis | N | N | N | N |
| ACE inhibitors | N | N | Y | N |
| AT1 blockers | Y | N | N | N |
| b-Blockers | Y | N | Y | Y |
| Dihydropyridines | Y | N | N | N |
| Diuretics | Y | N | Y | N |
| Nitrates | N | N | N | N |
| Lipid-lowering drugs | Y | N | Y | Y |

ACE, angiotensin-converting enzyme; AT, angiotensin receptor; ; BMI, body mass index; CAD, coronary artery disease; LVEF, left ventricular ejection fraction; MVD/AVD, mitral/aortic valve disease.

**Table S2.** AP Models’ parameters. Conductivities scalers.

|  | persAF | pAF | poAF | SR |
| --- | --- | --- | --- | --- |
| P_Na | 0.82 | 0.80 | 0.80 | 1.18 |
| g_Ca_L (Maleckar) | 0.15 | 0.01 | 0.01 | 0.61 |
| g_Ca_L (Tomek) | 0.92 | 0.30 | 0.01 | 0.55 |
| g_to | 0.14 | 0.20 | 0.79 | 1.21 |
| g_kur | 0.12 | 0.56 | 0.10 | 0.10 |
| g_K1 | 0.75 | 0.17 | 0.41 | 0.19 |
| g_Kr | 2.29 | 4.35 | 2.09 | 2.62 |
| g_Ks | 5.59 | 5.17 | 0.58 | 7.32 |
| g_b_Na | 2.74 | 0.18 | 0.66 | 0.22 |
| g_b_Ca | 0.54 | 0.26 | 0.11 | 0.11 |
| i_NaK_max | 0.35 | 0.28 | 0.28 | 0.61 |
| i_CaP_max | 9.49 | 1.24 | 0.17 | 0.13 |
| K_NaCa | 1.18 | 0.15 | 0.36 | 0.13 |
| alpha_rel (RyR) | 0.10 | 0.11 | 0.25 | 0.13 |
| I_up_max (SERCA) | 1.58 | 9.63 | 0.89 | 9.80 |

**Table S3.** Segmentation metrics for the synthetic dataset. Mean, standard deviation, 25%, 50%, 75% quantiles are calculated for accuracy, IoU (Intersection over Union), segment offset, and onset.

|  | Train (n = 474) |  |  |  | Train (n = 117) |  |  |  |
| --- | --- | --- | --- | --- | --- | --- | --- | --- |
|  | IoU,<br>% | Offset,<br>ms | Onset,<br>ms | Accuracy,<br>% | IoU,<br>% | Offset,<br>ms | Onset,<br>ms | Accuracy,<br>% |
| **mean** | 93 | 58 | -38 | 95 | 93 | 45 | -40 | 95 |
| **std** | 8 | 238 | 77 | 5 | 6 | 181 | 58 | 5 |
| **25%** | 91 | 0 | -6 | 93 | 91 | 0 | -60 | 95 |
| **50%** | 95 | 20 | -2 | 96 | 95 | 20 | -20 | 97 |
| **75%** | 97 | 60 | 0 | 98 | 97 | 50 | 0 | 98 |

**Table S4.** AP markers, experiment vs model.

|  |  | APD |  | Resting potential |  | AP Amplitude |  |
| --- | --- | --- | --- | --- | --- | --- | --- |
|  | CL | experim | model | experim | model | experim | model |
| persAF | 344 | 65 | 64 | -75.68 | -75.39 | 118.71 | 112.46 |
|  | 516 | 69 | 70 | -77.82 | -77.22 | 118.71 | 115.88 |
|  | 1032 | 74 | 76 | -79.80 | -80.12 | 121.92 | 119.69 |
|  | 2064 | 81 | 81 | -81.18 | -78.79 | 123.90 | 119.16 |
| pAF | 206 | 75 | 78 | -71.72 | -71.61 | 104.68 | 103.05 |
|  | 258 | 81 | 83 | -75.84 | -75.01 | 113.37 | 110.35 |
|  | 344 | 93 | 89 | -76.14 | -74.96 | 116.42 | 113.58 |
|  | 516 | 92 | 101 | -81.63 | -82.35 | 127.11 | 123.40 |
| poAF | 344 | 95 | 92 | -78.28 | -76.55 | 107.27 | 111.54 |
|  | 516 | 103 | 109 | -79.50 | -79.47 | 116.73 | 117.17 |
|  | 1032 | 132 | 136 | -81.63 | -81.23 | 124.05 | 120.85 |
|  | 2064 | 143 | 153 | -82.09 | -81.93 | 124.21 | 122.55 |
| SR | 344 | 125 | 114 | -82.40 | -78.45 | 133.06 | 123.08 |
|  | 516 | 147 | 143 | -82.40 | -80.25 | 133.36 | 128.75 |
|  | 1032 | 187 | 199 | -87.13 | -86.43 | 142.97 | 137.83 |
|  | 2064 | 221 | 220 | -85.45 | -82.84 | 141.60 | 137.00 |

**Figure S1.**
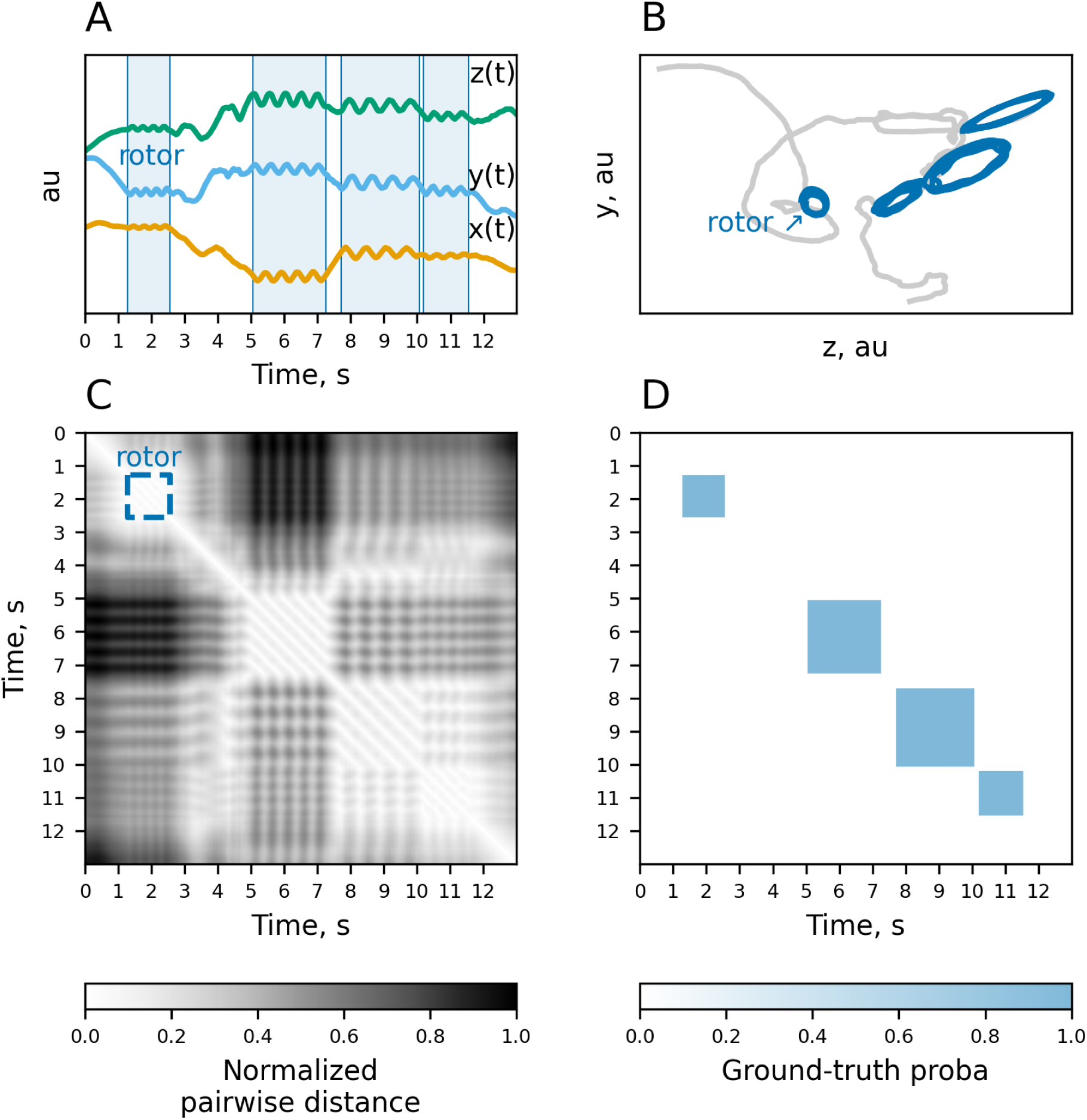
Example of the synthetic trajectory. (A) Trajectory coordinates. Blue shaded regions correspond to rotor segments. (B) Trajectory projection onto zy plane. Dark-blue and light-grey parts are rotor and non-rotor segments, respectively. (C) Normalized pairwise distances (euclidean) between trajectory points. This matrix is used as input for the neural network. (D) Ground-truth probability of the rotor (𝑃). If *i^th^* and *j^th^* trajectory points belong to the same rotor segment then *P*[*i, j*] and *P*[*j, i*] are supposed to be equal to 1(0, otherwise).

**Figure S2.**
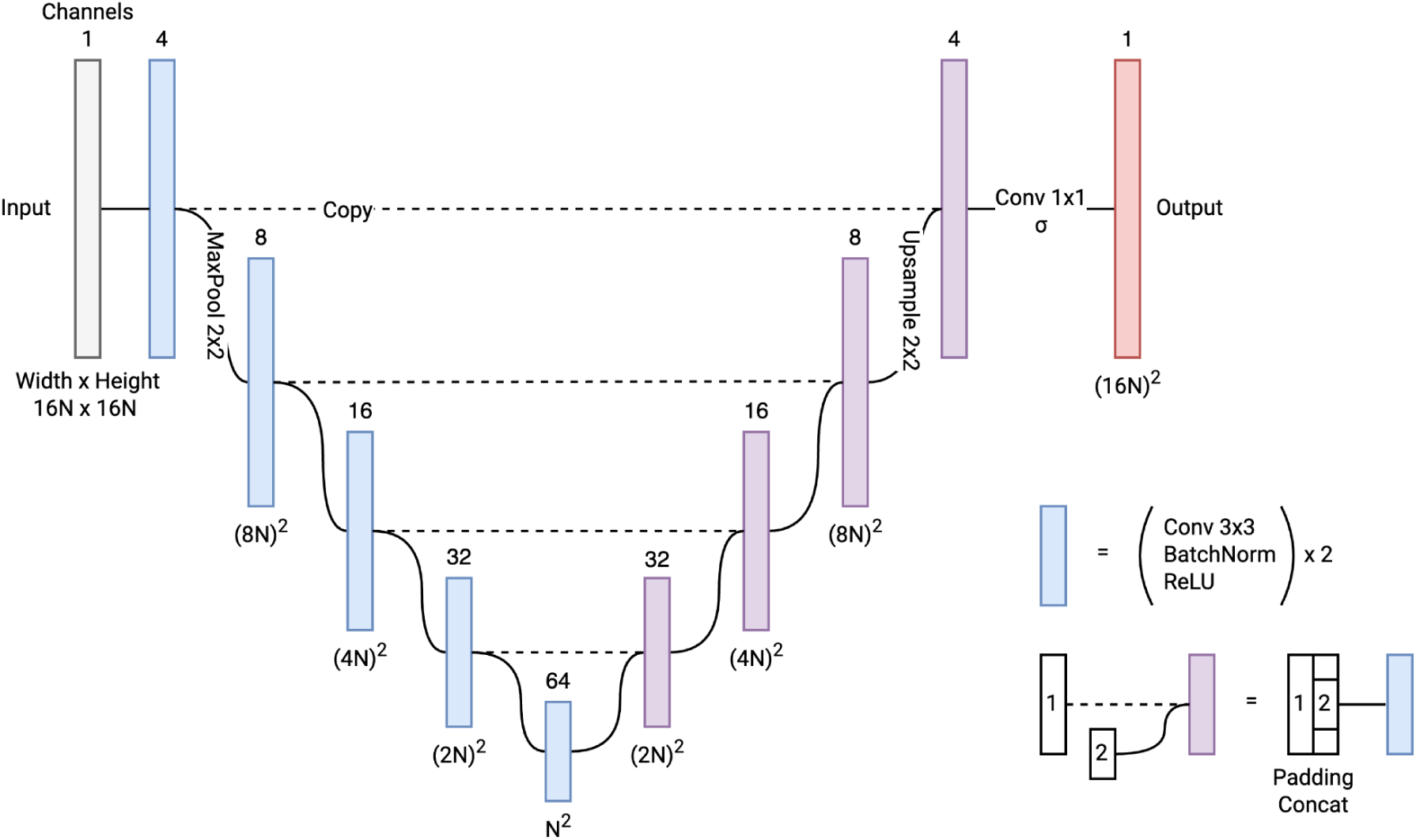
Neural network architecture.

**Figure S3.**
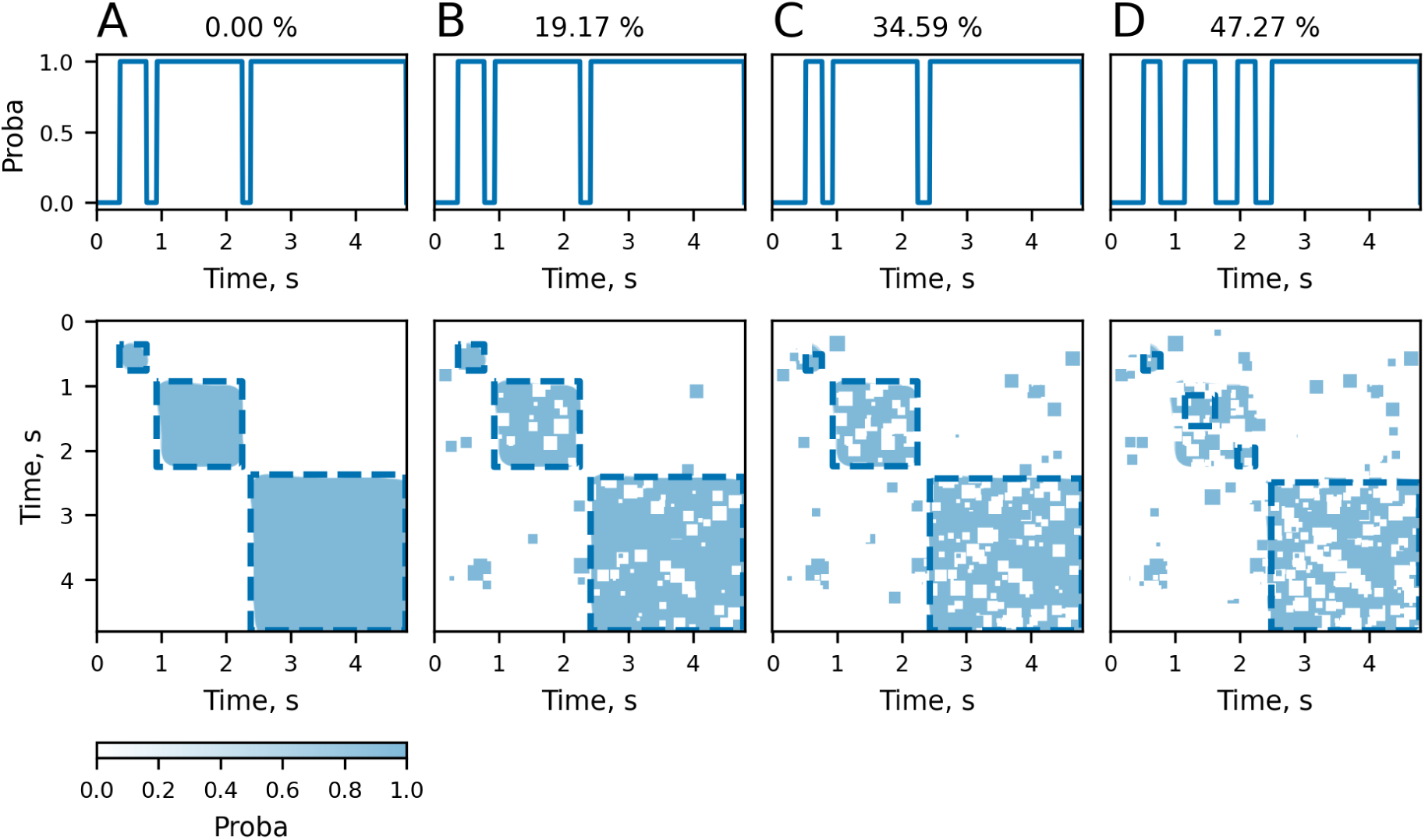
Segmentation of 2D matrices is robust. Example of the segmentation result (A). Panels B, C, and D correspond to the progressive increase in the misclassification rate (% of the misclassified pixels given in the subtitle). Even with 35% of the misclassified pixels, three reentrant driver segments were identified correctly.

**Figure S4.**
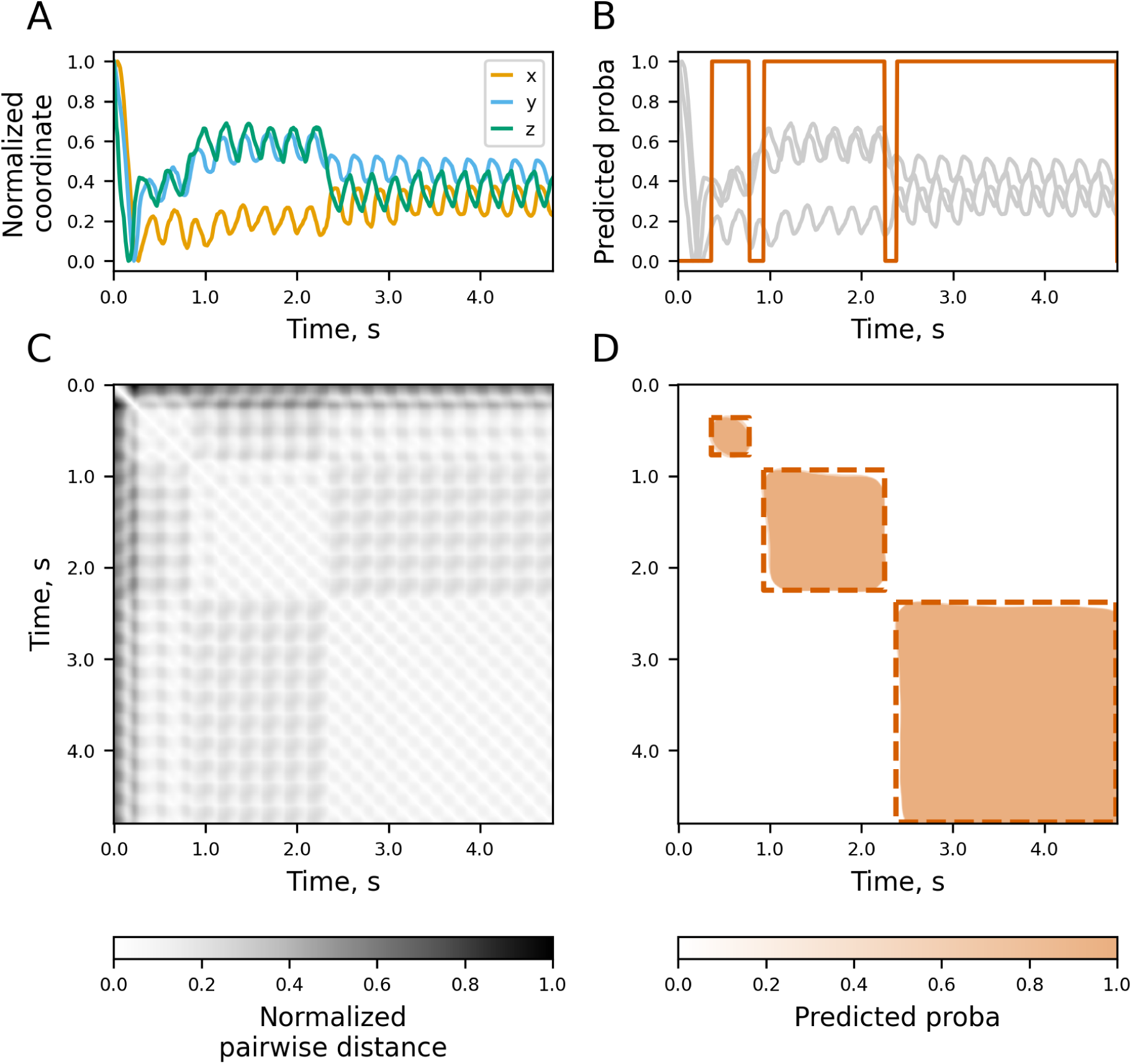
Example of the segmentation of the actual trajectory from the whole-atria simulations. (A) Trajectory coordinates. (B) Predicted probability (orange) of trajectory segment corresponding to reentrant driver. (C) Pairwise normalized distance (euclidean) between trajectory points. This matrix is used as an input to the convolutional neural network. (D) Predicted probability of the reentrant activity for the input 2D matrix. Three square-like orange blocks were classified as corresponding to reentrant drivers, dashed square section is used for final segmentation shown in (B).

**Figure S5.**
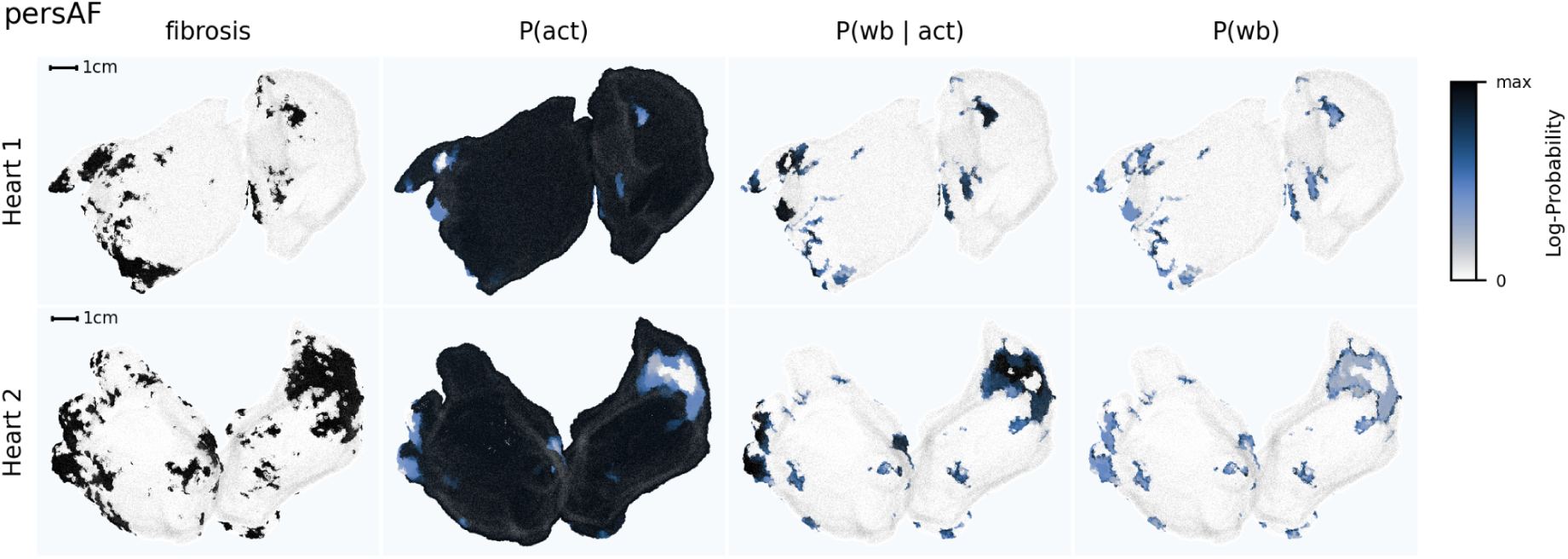
Example of activation/wavebreak probabilities for the persAF. The first column corresponds to the fibrosis distribution (white and black are normal and fibrotic regions, respectively). The following columns from left to right are activation probability 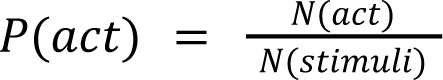, conditional wavebreak probability given 𝑁(𝑠𝑡𝑖𝑚𝑢𝑙𝑖) the activation 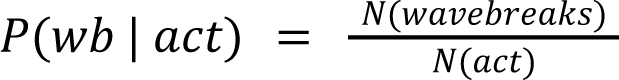, unconditional wavebreak probability 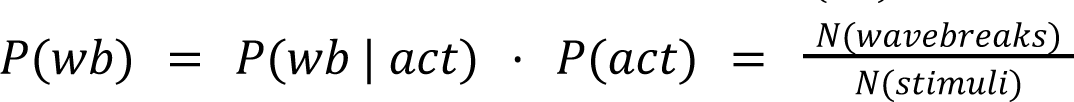. These probabilities were calculated over the first *2.5 s* of the simulation when the tissue was activated by pacing (not by arrhythmogenic reentrant drivers).

**Figure S6.**
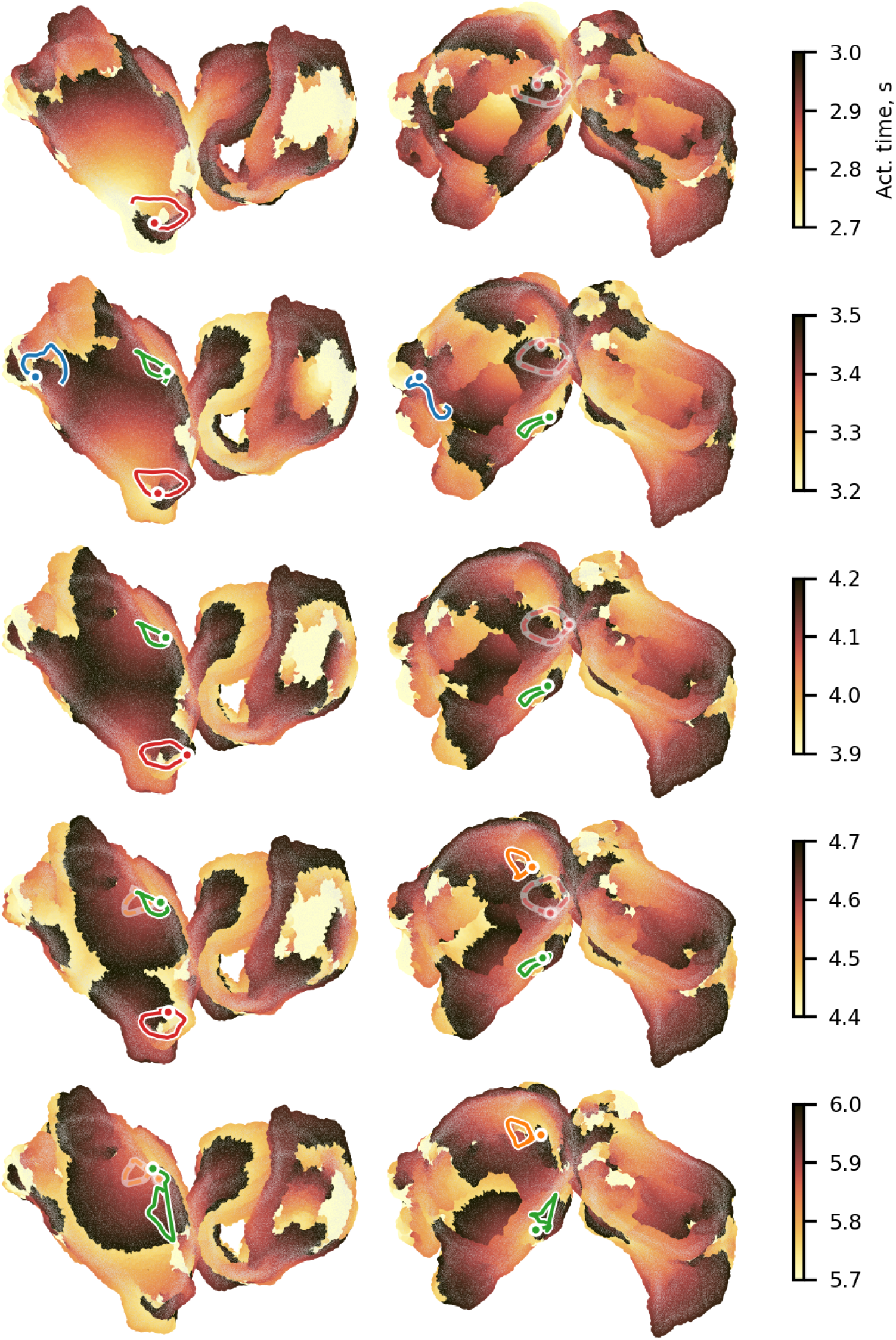
Activation sequence. Representative evolution of propagation pattern. Arrhythmogenic pattern is driven by several simultaneous reentrant drivers shown by colored curves.

**Figure S7.**
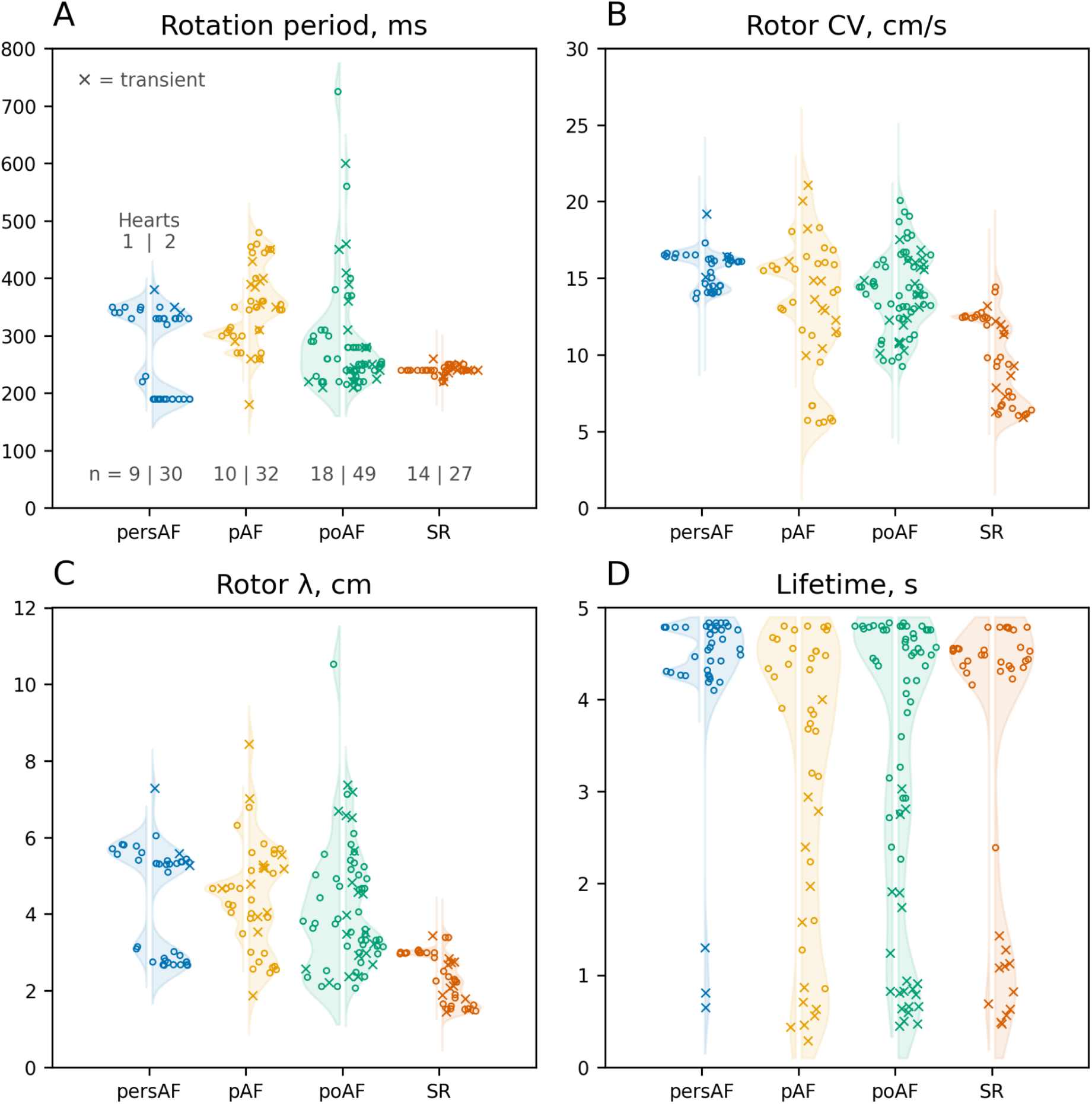
Reentrant drivers characteristics. Rotation period (A), conduction velocity (B), wavelength (C), and lifetime (D). ‘x’ marks transient rotors, *i.e.* it self-terminates before the end of the simulation. Multimodality is caused by reentrant drivers located in different anatomical regions. For example, the high CL mode of the persAF group corresponds to rotors around pulmonary veins (anatomical block).

**Figure S8.**
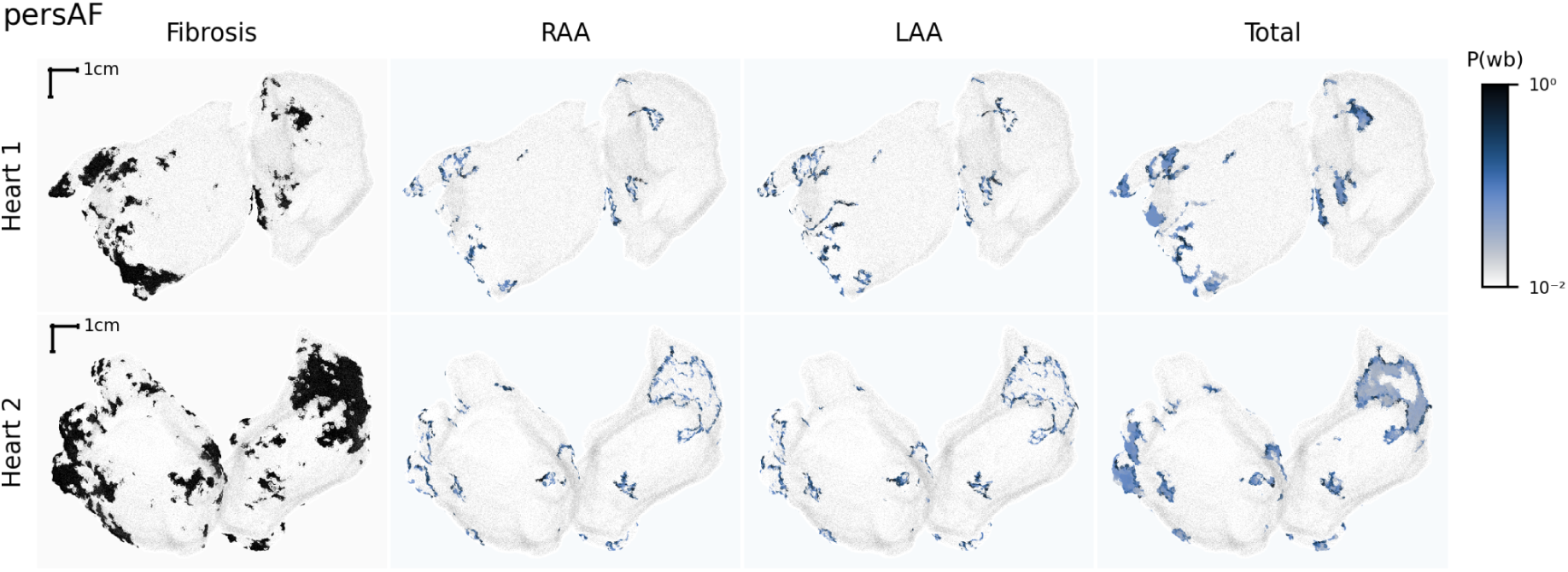
Wavebreak probability, P(wb), is independent of the pacing site. The first column is the fibrosis distribution (white and black are normal and fibrotic regions, respectively). The second and third columns correspond to the two simulations where stimuli were delivered to RAA and LAA, respectively. The last column shows the P(wb) averaged over all 40 simulations (pacing sites). The spatial distribution of the P(wb)depends on the fibrosis distribution and very similar across different simulations.

